# ICA-based Denoising Strategies in Breath-Hold Induced Cerebrovascular Reactivity Mapping with Multi Echo BOLD fMRI

**DOI:** 10.1101/2020.08.18.256479

**Authors:** Stefano Moia, Maite Termenon, Eneko Uruñuela, Gang Chen, Rachael C. Stickland, Molly G. Bright, César Caballero-Gaudes

## Abstract

Performing a BOLD functional MRI (fMRI) acquisition during breath-hold (BH) tasks is a non-invasive, robust method to estimate cerebrovascular reactivity (CVR). However, movement and breathing-related artefacts caused by the BH can substantially hinder CVR estimates due to their high temporal collinearity with the effect of interest, and attention has to be paid when choosing which analysis model should be applied to the data. In this study, we evaluate the performance of multiple analysis strategies based on lagged general linear models applied on multi-echo BOLD fMRI data, acquired in ten subjects performing a BH task during ten sessions, to obtain subject-specific CVR and haemodynamic lag estimates. The evaluated approaches range from conventional regression models including drifts and motion timecourses as nuisance regressors applied on single-echo or optimally-combined data, to more complex models including regressors obtained from multi-echo independent component analysis with different grades of orthogonalization in order to preserve the effect of interest, i.e. the CVR. We compare these models in terms of their ability to make signal intensity changes independent from motion, as well as the reliability as measured by voxelwise intraclass correlation coefficients of both CVR and lag maps over time. Our results reveal that a conservative independent component analysis model applied on the optimally-combined multi-echo fMRI signal offers the largest reduction of motion-related effects in the signal, while yielding reliable CVR amplitude and lag estimates, although a conventional regression model applied on the optimally-combined data results in similar estimates. This work demonstrate the usefulness of multi-echo based fMRI acquisitions and independent component analysis denoising for precision mapping of CVR in single subjects based on BH paradigms, fostering its potential as a clinically-viable neuroimaging tool for individual patients. It also proves that the way in which data-driven regressors should be incorporated in the analysis model is not straight-forward due to their complex interaction with the BH-induced BOLD response.

## 1 Introduction

Cerebrovascular reactivity (CVR) is a physiological response of the cerebral vessels to vasodilatory or vasoconstrictive stimuli. Mapping of the CVR response provides an important indicator of cerebrovascular health. In recent years, functional magnetic resonance imaging (fMRI), either based on the blood oxygenation level-dependent (BOLD) contrast, arterial spin labelling, or a mixture of both, has demonstrated its effectiveness as a method to assess CVR. As a result, its use is spreading into clinical practice, where its potential as a diagnostic measure is being ascertained in different diseases, spanning from vascular diseases (Hartkamp, Bokkers, van Osch, de Borst, & Hendrikse, 2017; Markus & Cullinane, 2001; Webster et al., 1995; Ziyeh et al., 2005), to stroke and aphasia (Krainik, Hund-Georgiadis, Zysset, & Von Cramon, 2005; Van Oers et al., 2018), brain tumors (Fierstra et al., 2018; Zacà, Jovicich, Nadar, Voyvodic, & Pillai, 2014), neurodegenerative diseases (Camargo et al., 2015; Glodzik, Randall, Rusinek, & de Leon, 2013; Marshall et al., 2014), hypertension (Iadecola & Davisson, 2008; Leoni et al., 2011; Tchistiakova, Anderson, Greenwood, & Macintosh, 2014), lifestyle habits (Friedman et al., 2008; Gonzales et al., 2014), sleep apnea (Buterbaugh et al., 2015; Prilipko, Huynh, Thomason, Kushida, & Guilleminault, 2014), and traumatic brain injury or concussions (Churchill, Hutchison, Graham, & Schweizer, 2020; Markus & Cullinane, 2001).

CVR measurements are obtained by evoking a vasodilatory response during imaging. This is typically done by injecting intravenous acetazolamide, or by exposing the subject to gas challenges with computerised dynamic deployment of CO_2_ and O_2_. However, acetazolamide is an invasive technique not indicated for vulnerable subjects (e.g. elderly or children), while gas challenges require dedicated setups and can also cause discomfort in some subjects, which might induce anxiety and thus potentially bias CVR measurement (Urback, MacIntosh, & Goldstein, 2017). A valid alternative is the use of voluntary respiratory challenges, such as paced deep breathing or breath-hold (BH) tasks (Bright, Bulte, Jezzard, & Duyn, 2009; Kastrup, Li, Takahashi, Glover, & Moseley, 1998). In fact, it has been shown that CO_2_ changes in the blood due to breathing tasks elicit a CVR response that is equivalent to that of inhaled CO_2_ (Kastrup, Krüger, Neumann-Haefelin, & Moseley, 2001; Tancredi & Hoge, 2013). A BH task can be successfully implemented in young children and elderly subjects (Handwerker, Gazzaley, Inglis, & D’Esposito, 2007; Thomason, Burrows, Gabrieli, & Glover, 2005), and it is a robust measurement even if subjects are not able to hold their breath for as long as instructed (Bright & Murphy, 2013a). Moreover, BH-induced CVR has been shown to be a reliable measurement across different sessions of MRI, both in the short (same day) and long term (Peng, Yang, Chen, & Shih, 2019), in terms of spatial reliability (i.e. comparing variability of voxels across multiple sessions in one subject) and general reliability (i.e. average CVR value across sessions and within subjects) (Lipp, Murphy, Caseras, & Wise, 2015; Magon et al., 2009). Both short and long term reliability of BH-induced CVR were found to be comparable to that of other non-invasive means of estimating CVR, such as resting state fMRI (P. Liu et al., 2017), inhaled gas challenges (Dengel et al., 2017; Evanoff et al., 2020; Leung, Kim, & Kassner, 2016), Fourier modelling of a BH task (Pinto, Jorge, Sousa, Vilela, & Figueiredo, 2016), and a paced deep breathing task (Sousa, Vilela, & Figueiredo, 2014).

However, BOLD fMRI data exhibit signal variation arising from different sources, most of which corresponds to hardware-related artefacts and drifts, head motion, confounding physiological fluctuations, and other sources of noise (Bianciardi et al., 2009; Jorge, Figueiredo, van der Zwaag, & Marques, 2013). It is important that the signal variance associated with these confounding signals is accounted for and minimized during preprocessing or data analyses (Caballero-Gaudes & Reynolds, 2017; T. T. Liu, 2016). The artefacts induced by voluntary and involuntary movement are a particularly problematic source of noise for task-based fMRI experiments, mainly in block designs where large head movement leads to bias in estimates of the task-induced activity (Johnstone et al., 2006) and in particular experimental paradigms, such as in overt speech production where the articulation of words makes head movement considerable (Barch et al., 1999; Soltysik & Hyde, 2006; Xu et al., 2014). This concern with task-induced movement artefacts extends to respiration tasks: the experimental design is similar to that of block designs, but the amount of motion associated with paced breathing, deep breaths, or “recovery” breaths following a BH task can be very prominent and concur with the pattern of the task. Moreover, respiration can perturb the B0 field due to the change of air in the lungs (Raj, Anderson, & Gore, 2001) and introduce aliasing artefacts or pseudo-movement effects in the signal (Gratton et al., 2020; Pais-Roldán, Biswal, Scheffler, & Yu, 2018; Power, Lynch, et al., 2019).

There are different ways to account for motion effects on task-based fMRI data analysis. For instance, such effects can be reduced during acquisition by implementing an event-related task paradigm (Birn, Bandettini, Cox, & Shaker, 1999; Birn, Cox, & Bandettini, 2004). However, in a BH task the periods of apnoea are typically between 10 and 20 seconds in duration to achieve a robust and reproducible vasodilatory response (Bright & Murphy, 2013a; Magon et al., 2009), and are not readily adapted to a brief event-related design. The most straight-forward approach is then to include the realignment translation and rotation parameters in the analysis model (Friston, Williams, Howard, Frackowiak, & Turner, 1996). Including such timecourses, as well as their derivatives and non-linear expansions, in the regression model can account for part of the motion-related variance of the signal, thus improving the estimation of the task effects. Another widely adopted method to remove motion-related effects, as well as noise in general, is to decompose the fMRI data using Principal Component Analysis or Independent Component Analysis (ICA) in order to identify and remove components that are mostly related to motion or other sources of noise (Behzadi, Restom, Liau, & Liu, 2007; Griffanti et al., 2014; Muschelli et al., 2014; R. H. R. Pruim, Mennes, Buitelaar, & Beckmann, 2015; R. H. R. Pruim, Mennes, Rooij, et al., 2015; Salimi-Khorshidi, Smith, & Nichols, 2011).

Alternatively, noise in fMRI can be reduced by using multi-echo (ME) acquisitions that sample the data at multiple successive echo times (TE). A weighted combination of the multiple echoes based on each voxel’s 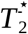 value (Posse et al., 1999) or temporal signal-to-noise ratio (Poser, Versluis, Hoogduin, & Norris, 2006) can smear out random noise and enhance the sensitivity to the BOLD contrast. In fact, compared with single-echo data, this optimal combination can improve the mapping of neuronal activity at 3 Tesla (Fernandez, Leuchs, Sämann, Czisch, & Spoormaker, 2017) and 7T (Puckett et al., 2018), with results comparable to other preprocessing techniques requiring extra data such as RETROICOR (Atwi et al., 2018). Optimal combination of multiple echo volumes can also improve BH-induced CVR mapping sensitivity, specificity, repeatability and reliability (Cohen & Wang, 2019).

Furthermore, assuming monoexponential decay, the voxelwise fMRI signal (S) can be expressed in signal percentage change as:

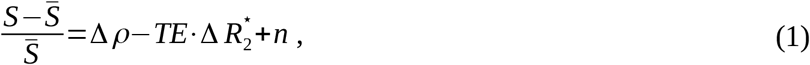

where Δ*ρ* represents non-BOLD related changes in net magnetisation, 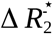 represents BOLD-related susceptibility changes, and *n* denotes the random noise (Kundu, Inati, Evans, Luh, & Bandettini, 2012). As the BOLD-related signal can be expressed as a function of the TE, whereas noise-related non-BOLD changes in the net magnetization are independent of TE, the information available in multiple echoes can be leveraged for the purpose of denoising. For example, in a dualecho acquisition where the first TE is sufficiently short, the first echo signal mainly captures changes in Δ*ρ* rather than in 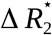. It is then possible to remove artefactual effects, through voxelwise regression, from the second echo signal acquired at a longer TE with appropriate BOLD contrast (Bright & Murphy, 2013b). Collecting more echoes opens up the possibility of applying ICA and classifying independent components into BOLD-related (i.e. describing 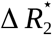 fluctuations with a linear TE-dependency) or noise (i.e. independent of TE, related to non-BOLD fluctuations in the net magnetization Δ*ρ*), an approach known as multi-echo independent component analysis (ME-ICA) (Kundu et al., 2013, 2012, 2017). Compared to single-echo data denoising, ME-ICA can improve the mapping of task-induced activation (DuPre, Luh, & Spreng, 2016; Gonzalez-Castillo et al., 2016; Lombardo et al., 2016), for example in challenging paradigms with slow-varying stimuli (Evans, Kundu, Horovitz, & Bandettini, 2015) or language mapping and laterality (Amemiya, Yamashita, Takao, & Abe, 2019). It also outperforms single-echo ICA-based denoising of restingstate fMRI data (Dipasquale et al., 2017), which is particularly beneficial more efficient and reliable functional connectivity mapping in individual subjects (Lynch, Power, Dubin, Gunning, & Liston, 2020) and in brain regions where traditional single-echo acquisitions offer reduced signal-to-noise ratio, such as the basal forebrain (Markello, Spreng, Luh, Anderson, & De Rosa, 2018). Furthermore, ME-ICA also enhances the deconvolution of neuronal-related signal changes (Caballero-gaudes, Moia, Panwar, Bandettini, & Gonzalez-castillo, 2019).

However, up to now, the operation of ME-ICA has not been evaluated thoroughly in experimental paradigms with unavoidable task-correlated artefacts. Under such scenarios, one question that remains open is how to obtain the right trade-off between removing as much noise as possible and saving as much signal of interest as possible (Bright & Murphy, 2015; Griffanti et al., 2014). In this study, we acquire ME-fMRI data during a BH task in 10 subjects acquired weekly over the course of 2.5 months, i.e. adopting a similar framework to recent precision functional mapping experiments (Gordon et al., 2017; Greene et al., 2020; Laumann et al., 2015; Lynch & Liston, 2020; Marek et al., 2018), and assess the efficiency of different nuisance regression models to remove artefacts that are highly correlated with the effect of interest, i.e. the CVR response. In particular, we compare traditional nuisance regression approaches, applied to single- or multi-echo data, and three different ME-ICA denoising approaches ranging from aggressive to conservative. For each denoising strategy, we assess the correlation of the cleaned signal with measures of motion, and evaluate the amplitude and lag of the CVR signal response in terms of their physiological interpretability and inter-session reliability.

## 2 Material and methods

### 2.1 Participants

Ten healthy subjects with no record of psychiatric or neurological disorders (5F, age range 24-40 y at the start of the study) underwent ten MRI sessions in a 3T Siemens PrismaFit scanner with a 64-channel head coil. Each session took place one week apart, on the same day of the week and at the same time of the day.

All participants had to meet several further requirements, i.e. being non-smokers and refrain from smoking for the whole duration of the experiment, and not suffering from respiratory or cardiac health issues. They were also instructed to refrain from consuming caffeinated drinks for two hours before the session. Informed consent was obtained before each session, and the study was approved by the local ethics committee.

### 2.2 Data acquisition and MRI session

Within the MRI session, subjects performed a BH task while T2*-weighted ME-fMRI data was acquired with the simultaneous multislice (a.k.a. multiband, MB) gradient-echo planar imaging sequence provided by the Center for Magnetic Resonance Research (CMRR, Minnesota) (Moeller et al., 2010; Setsompop et al., 2012) with the following parameters: 340 scans, TR = 1.5 s, TEs = 10.6/28.69/46.78/64.87/82.96 ms, flip angle = 70°, MB acceleration factor = 4, GRAPPA = 2 with Gradient-echo reference scan, 52 slices with interleaved acquisition, Partial-Fourier = 6/8, FoV = 211×211 mm^2^, voxel size = 2.4×2.4×3 mm^3^, Phase Encoding = AP, bandwidth=2470 Hz/px, LeakBlock kernel reconstruction (Cauley, Polimeni, Bhat, Wald, & Setsompop, 2014) and SENSE coil combination (Sotiropoulos et al., 2013). Single-band reference (SBRef) images were also acquired for each TE. The BH task was preceded by 64 minutes of ME-fMRI scanning, consisting of three task-based and four 10-minute resting state acquisitions, which are not part of the current study. The BH task always followed a resting state run. A pair of Spin Echo echo planar images (EPI) with opposite phase-encoding (AP or PA) directions and identical volume layout (TR = 2920 ms, TE = 28.6 ms, flip angle = 70°) were also acquired before each functional run in order to estimate field distortions, similarly to the Human Connectome Project protocol (Glasser et al., 2016). A T1-weighted MP2RAGE image (Marques et al., 2009) (TR = 5 s, TE = 2.98 ms, TI1 = 700 ms, TI2 = 2.5 s, flip angle 1 = 4°, flip angle 2 = 5°, GRAPPA = 3, 176 slices, FoV read = 256 mm, voxel size = 1×1×1 mm3, TA = 662 s) and a T2-weighted Turbo Spin Echo image (Hennig, Nauerth, & Friedburg, 1986) (TR = 3.39 s, TE = 389 ms, GRAPPA = 2, 176 slices, FoV read = 256 mm, voxel size = 1×1×1 mm3, TA = 300 s) were also collected at the end and at the beginning of each MRI session, respectively.

During the fMRI acquisition runs exhaled CO_2_ and O_2_ levels were monitored and recorded using a nasal cannula (Intersurgical) with an ADInstruments ML206 gas analyser unit and transferred to a BIOPAC MP150 physiological monitoring system where scan triggers were simultaneously recorded. Photoplethismography and respiration effort data were also measured via the BIOPAC system, but these physiological signals were not used in the current study. All signals were sampled at 10 kHz.

### 2.3 Breath-hold task

Following Bright and Murphy (2013a), the BH paradigm consisted of eight repetitions of a BH trial composed of four paced breathing cycles of 6 s each, an apnoea (BH) of 20 s, an exhalation of 3 s, and 11 s of “recovery” breathing (unpaced) (i.e. total trial duration of 58 s) (Figure 1). Subjects were instructed prior to scanning about the importance of the exhalations preceding and following the apnoea. Without these exhalations providing CO_2_ measurements, the change in systemic CO_2_ levels achieved by each BH cannot be robustly estimated; as a result, CVR (%BOLD/mmHg CO_2_ change) cannot be estimated quantitatively. Participants were instructed textually throughout the task through a mirror screen located in the head coil.

**Figure 1:**
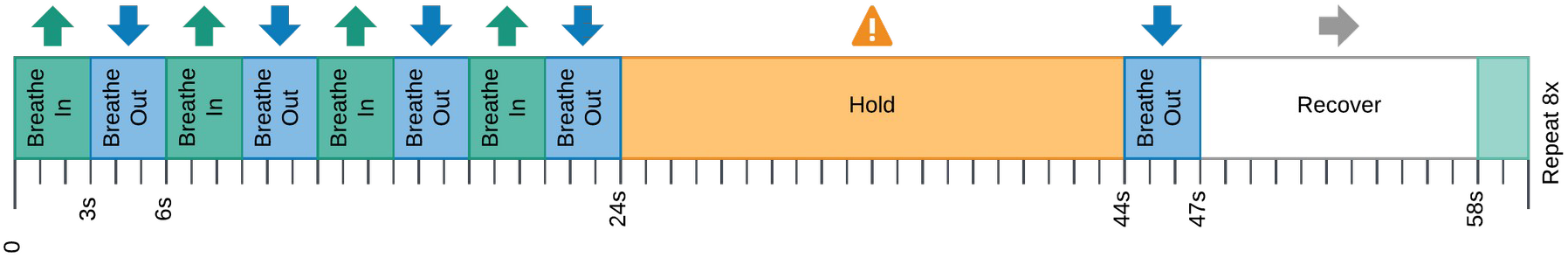
Schematic of Breath-Hold trial. Apnoea was preceded and followed by exhalations.

### 2.4 MRI data preprocessing

The DICOM files of the MRI data were transformed into nifti files with dcm2nii (Li, Morgan, Ashburner, Smith, & Rorden, 2016) and formatted into Brain Imaging Data Structure (Gorgolewski et al., 2016) with heudiconv (Halchenko et al., 2019).

MRI data were minimally preprocessed with custom scripts based mainly in FSL (Jenkinson, Beckmann, Behrens, Woolrich, & Smith, 2012), AFNI (Cox, 1996), and ANTs (Tustison et al., 2014). In brief, the T2-weighted image was skull-stripped and co-registered to the MP2RAGE image along with the brain mask. The latter was applied to the MP2RAGE image, that was then segmented into gray matter (GM), white matter (WM) and cerebrospinal fluid tissues (Avants, Tustison, Wu, Cook, & Gee, 2011). The MP2RAGE image was normalised to an asymmetric version of the MNI152 6^th^ generation template at 1 mm resolution (Grabner et al., 2006), while the T2-weighted volume was co-registered to the skull-stripped single-band reference image (SBRef) of the first echo. The first 10 volumes of the functional data were discarded to allow the signal to achieve a steady state of magnetisation. Image realignment to the SBRef was computed on the first echo, and the estimated rigid-body spatial transformation was then applied to all other echoes (Jenkinson, Bannister, Brady, & Smith, 2002; Jenkinson & Smith, 2001). A brain mask obtained from the SBRef volume was applied to all the echoes. The different echo timeseries were optimally combined (OC) voxelwise by weighting each timeseries contribution by its *T*_2_ value (Posse et al., 1999). Next, ME-ICA decomposition was performed on each run independently with tedana (DuPre et al., 2019) using the minimum description length criterion for estimation of the number of components (Harris, 1978; Li et al., 2016). The independent components (ICs) were then manually classified by SM and CCG into two categories (rejected or accepted components) based on temporal, spatial, spectral and TE-dependence features of each component (Griffanti et al., 2017). The manual classifications are available in the data repository. A distortion field correction was performed on the OC volume with Topup (Andersson, Skare, & Ashburner, 2003), using the pair of spin-echo EPI images with reversed phase encoding acquired before the ME-EPI acquisition (Glasser et al., 2016). Finally, the BOLD timeseries was converted in signal percentage change. For comparison, the dataset acquired at the second echo time (TE_2_ = 28.6 ms) was used as a surrogate for standard single-echo (SE) acquisitions. This volume followed the same preprocessing steps as the OC volume, except for the optimal combination and the ICA decomposition.

### 2.5 CO_2_ trace processing and CVR estimation

The files exported from the AcqKnowledge software were transformed and formatted into BIDS with phys2bids (The phys2bids developers et al., 2019).

The CO_2_ timecourse was processed using custom scripts in Python 3.6.7. Briefly, the CO_2_ timecourse was downsampled to 40 Hz to reduce computational costs. The end-tidal peaks were automatically and manually individuated. The amplitude envelope was obtained by linearly interpolating between the end-tidal peaks and it was then demeaned and convolved with a canonical HRF to obtain the P_ET_CO_2_*hrf* trace. In order to account for measurement delay, the P_ET_CO_2_*hrf* trace was shifted to maximise the cross-correlation with the average timecourse of an eroded version of the GM mask (bulk shift) (Yezhuvath, Lewis-Amezcua, Varghese, Xiao, & Lu, 2009). This step was performed on both OC and the SE data (see Supplementary figure 1).

A general linear model (L-GLM) approach was adopted in this study for CVR estimation (Moia, Stickland, et al., 2020) in order to model temporal offsets between the P_ET_CO_2_ recording and the CVR response across voxels that occur due to measurement and physiological delays (Donahue et al., 2016; Geranmayeh, Wise, Leech, & Murphy, 2015; Murphy, Harris, & Wise, 2011; Sousa et al., 2014; Tong, Bergethon, & Frederick, 2011). Sixty shifted versions of the P_ET_CO_2_*hrf* trace were created, ranging between ±9 s from the bulk shift, with a shift increment of 0.3 s (fine shift). This temporal range was based on previous literature, which rarely reports haemodynamic lags over ±8 s in healthy individuals (Bright et al., 2009; Donahue et al., 2016; Sousa et al., 2014). For each shift, a GLM was defined with a design matrix comprised of the shifted P_ET_CO_2_*hrf* timecourse as the regressor of interest, and different combinations of nuisance regressors (see below) in order to examine their efficiency in modelling artefactual signals of the voxel timeseries that might degrade CVR estimates. The simultaneous fitting of the nuisance regressors and the regressor of interest (i.e. the shifted P_ET_CO_2_*hrf* trace) is preferable, rather than denoising via nuisance regression prior to the analysis (Jo et al., 2013; Lindquist, Geuter, Wager, & Caffo, 2019; Moia, Stickland, et al., 2020).

Five different modelling strategies were evaluated, varying which nuisance regressors were included in the design matrix or how they were derived from ME-ICA:

1. A L-GLM model on the SE data where the design matrix includes the motion parameters and their temporal derivatives (denoted as *Mot*), Legendre polynomials of up to the fourth order (denoted as *Poly*), together with the P_ET_CO_2_*hrf* trace (SE-MPR):

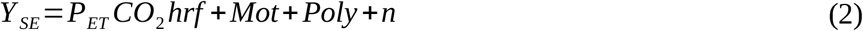
2. The same model applied on the OC data (OC-MPR):

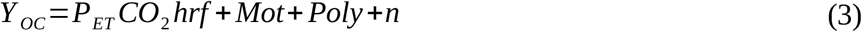
3. An *aggressive* model applied on the OC data (ME-AGG) in which the design matrix also includes the timecourses of the ME-ICA rejected components (denoted as *IC_rej_*) added to the design matrix of the L-GLM, orthogonalised with respect to the motion parameters, their temporal derivatives, and Legendre polynomials of up to the fourth order. This orthogonalisation step was performed to maintain a low Variance Inflation Factor in this model, and thus not bias the CVR estimation, without altering the relative variance explained by the original nuisance regressors and the regressor of interest (Mumford, Poline, & Poldrack, 2015):

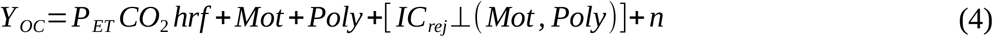
4. A *moderate* model applied on the OC data (ME-MOD) in which the timecourses of the ME-ICA rejected components are also orthogonalised with respect to the P_ET_CO_2_*hrf* trace (i.e. the regressor of interest describing the CVR response):

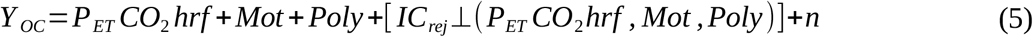
5. A *conservative* model applied on the OC data (ME-CON) in which the timeseries of the ME-ICA rejected components are orthogonalised with respect to the P_ET_CO_2_*hrf* trace and the ME-ICA accepted components (denoted as *IC_acc_*):

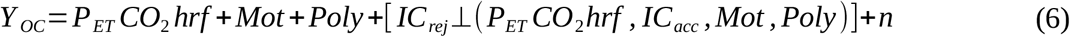

In the models above, *Y_SE_* and *Y_OC_* are the SE and OC voxel timeseries respectively and *n* denotes the random noise.

For each modelling strategy and each of the sixty shifted P_ET_CO_2_*hrf* traces, the corresponding L-GLM was fitted via orthogonal least squares using AFNI. Then, for each voxel, the beta coefficient (i.e. weight) of the best fine-shifted P_ET_CO_2_*hrf* trace, corresponding to the L-GLM model with maximum coefficient of determination (R^2^), was selected. Finally, the beta coefficients expressed in BOLD signal percentage change over Volts (BOLD_SPC_/V) were rescaled to be expressed in BOLD percentage over millimetres of mercury (%BOLD/mmHg) as indicated by the gas analyser manufacturer^1^.

In this way, a lag-optimised CVR map and a t-value map were obtained, together with the associated lag map representing the voxelwise delay from the bulk shift, for each analysis pipeline. To account for sixty comparisons computed in the L-GLM approach (one per regressor), the CVR and lag maps were thresholded at p<0.05 adjusted with the Šidák correction (Bright, Tench, & Murphy, 2017; Šidák, 1967), and the voxels that were not statistically significant were excluded. The maps were further thresholded on the basis of the lag: those voxels in which the optimal lag was at or adjacent to the boundary (i.e. |*lag*|≥8.7 *s*) were considered not truly optimised and not readily physiologically plausible in healthy subjects and therefore masked in all maps (Moia, Stickland, et al., 2020).

### 2.6 Evaluation of motion removal across denoising strategies

For each type of L-GLM analysis, 4-D volumes representing the modelled noise variance were reconstructed by multiplying the optimised beta coefficient maps of the nuisance regressors by their timeseries using 3dSynthesize in AFNI. Then, they were subtracted from the OC or the SE data to obtain five different denoised datasets. DVARS, the root of the spatial mean square of the first derivative of the signal (Smyser et al., 2010), was computed on each denoised dataset as:

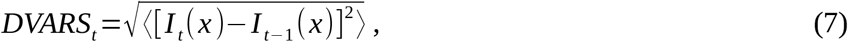

where *I_t_*(*x*) is the image intensity of voxel *x* and at time *t* and 〈···〉 indicates the spatial average over the whole brain. These DVARS timeseries were compared with the Framewise Displacement (FD) time courses (Power, Barnes, Snyder, Schlaggar, & Petersen, 2012), computed using the realignment parameters estimated during preprocessing using the fsl_motion_outliers tool as:

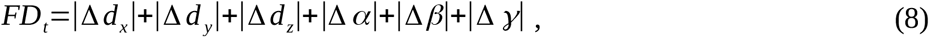

where *t* denotes the time, *d_x_, d_y_, d_z_* are the translational displacements along the three axes, *α, β, γ* are the rotational displacements of pitch, yaw, and roll, and Δ *d_x_*=*d*_*x,t*–1_–*d_x,t_* (and similarly for the other parameters). DVARS was also computed on the SE volume before preprocessing (SE-PRE) to serve as a reference, as its relationship with FD should be at its maximum prior to the effects of motion being removed.

In order to test the moderating effect of each analysis on the relationship between DVARS and FD, a Linear Mixed Effects (LME) model was set up using the lme4 and lmer packages (Bates, Mächler, Bolker, & Walker, 2015; Kuznetsova, Brockhoff, & Christensen, 2017) in R (R Core Team, 2020), computing the p value with Satterthwaite’s method (Satterthwaite, 1946), and accounting for the random effect of subject and session. The model was formulated as following in R equation notation:

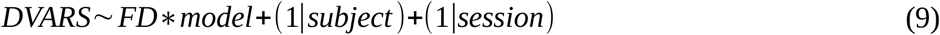

Then, the same model was also used to assess differences in motion removal between pairs of denoising strategies. The results were threshold at p=0.05 corrected with the Šidák correction (Šidák, 1967).

In order to visualise the CVR responses to a BH trial, the average timeseries within GM was extracted from each denoised dataset from each model SE-PRE, SE-MPR, OC-MPR, ME-AGG, ME-MOD, ME-CON, as well as from SE-PRE. These timeseries were transformed to BOLD percentage signal change, then the response to individual BH trials from each session were extracted using the timing of the third paced breathing cycle as a reference onset, and averaged together for each subject. The DVARS and FD timeseries followed the same process, except that the FD timeseries were not expressed in percentage.

Finally, the amount of BH trials necessary to achieve a robust estimation of the BH response was computed for each denoising approach. The Manhattan distance from a pool of a gradually increasing number of trials to the average BOLD response over all BH trials (across the ten sessions, 80 trials in total) was also computed for each analysis model and subject.

### 2.7 Comparison of CVR and lag estimation and reliability across denoising strategies

For each denoising strategy, the average CVR and lag values across the significant voxels in GM and WM was computed for all subjects and all sessions, in addition to the amount of statistically significant voxels in the thresholded CVR maps.

In order to compare the results of the different denoising strategies, the thresholded CVR and lag maps of each session were normalised with a nearest neighbour interpolation to the MNI152 template (Grabner et al., 2006). Then, a LME model was computed voxelwise using 3dLMEr (Chen, Saad, Britton, Pine, & Cox, 2013), considering the effect of subjects and sessions as random effects. The model was formulated as following in R equation notation:

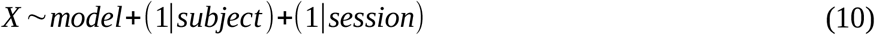

where *X* represents either the CVR or the lag value of each voxel. The same model was used to perform pairwise comparisons between the different strategies.

After normalising the t-value maps, the normalised CVR and lag maps were used to compute the intraclass correlation coefficient (ICC). ICC was computed voxelwise using a regularized multilevel mixed effect model in 3dICC (AFNI) in order to take into account the standard error of CVR and lag for each session in the ICC estimation (Chen et al., 2018). ICC assesses the reliability of a metric by comparing the intersubject, intrasubject, and total variability of that metric, which is equivalent to:

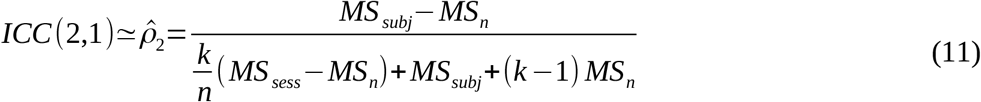

where *MS_subj_, MS_sess_*, and *MS_n_* are the mean squares of the effects of subjects, sessions, and residuals respectively, *k* = 10 is the number of sessions, and *n*= 7 the number of subjects (Chen et al., 2018; Mcgraw & Wong, 1996; Shrout & Fleiss, 1979). ICC(2,1) was chosen since both subjects and sessions were considered random effects. High ICC scores indicate high reliability, where the intrasubject variability is lower than the intersubject variability. Note that, since 3dICC uses the t-statistic map associated with the estimation of the CVR, CVR and lag maps used in this computation were thresholded only on the basis of the lag and not on the basis of the t-statistic.

### 2.8 Methods and data availability

In order to guarantee the replication of methods and results, all the code has been prepared to be run in a Singularity container based on a Ubuntu 18.04 Neurodebian OS. The container is publicly available at https://git.bcbl.eu/smoia/euskalibur_container, the methods pipeline is available at https://github.com/smoia/EuskalIBUR_dataproc, while all of the MRI images, physiological data, and manual classification used in this study will be available in OpenNeuro (EuskalIBUR dataset).

## 3 Results

Three subjects were excluded due poor performance of the BH task in part of the sessions, mainly due to inadequate execution of the exhalations preceding and following the apnoea which prevented accurate determination of the P_ET_CO_2_*hrf* traces. These traces are shown in red in Figure 2 that plots the PetCO_2_hrf trace for all subjects and sessions. Hence, only the seven subjects that had all ten session were used for subsequent analyses (4F, age 25-40y).

**Figure 2:**
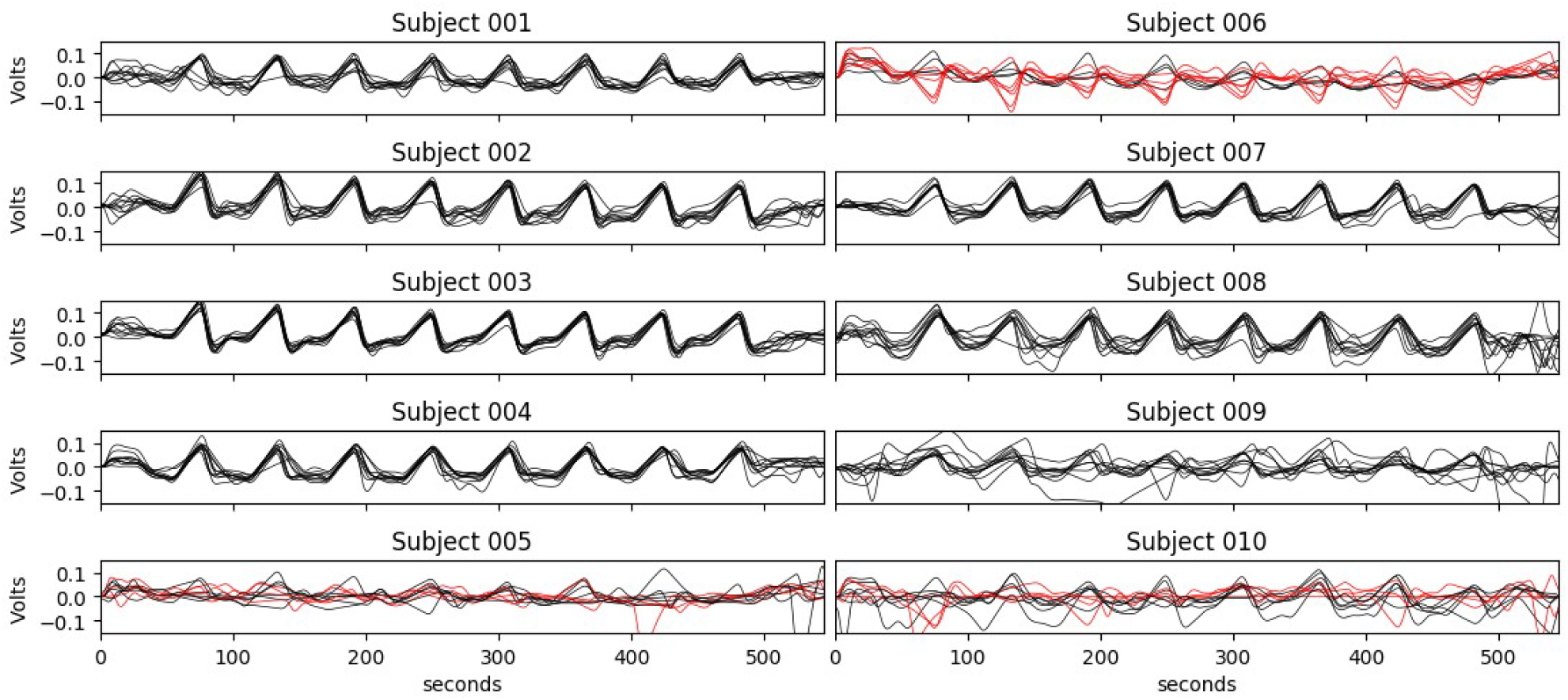
P_ET_CO_2_hrf trace for all subjects and all sessions. Rejected sessions are plotted in red. Rejection was based on having less than three proper P_ET_CO_2_ increases after breathholds or having more P_ET_CO_2_ decreases than increases after breathholds. Note that the first session of subject 10 was lost due to a software malfunction during acquisition.

### 3.1 Evaluation of motion removal across denoising strategies

Figure 3a illustrates the relationship between FD and DVARS in the raw data (SE-PRE) and after removing the reconstructed noise of each analysis model from the SE or OC volume for a representative subject; each point represents a timepoint and each line represents the linear regression between both timeseries in one session. The corresponding figures for the remaining subjects are available as Supplementary Material (supplementary figure 2). Figure 3b shows the same plot considering all the subjects and sessions. The modulating effect of the denoising approaches on the relation between DVARS and FD was tested with a LME model that was found to be significant (F(6,161181)=34597, p<0.001). To further investigate the significant differences between analysis strategies, Table 1 reports the results of the same LME model considering pairwise combinations of all of the denoising approaches. From both Figure 3 and Table 2, it can be seen the optimal combination (OC-MPR) of ME data reduces DVARS compared to single-echo (SE-MPR). Although a similar relationship is observed between DVARS and FD in both approaches, OC-MPR significantly reduces the impact of FD compared to SE-MPR (*β* =715.10, CI95 [710.17, 720.04], p<0.001). This relationship is even more mitigated in the moderate (ME-MOD) (*β* =145.40, CI_95_ [141.92, 148.88], p<0.001) and conservative (ME-CON) (*β* =146.69, CI_95_ [143.05, 150.33], p<0.001) denoising approaches, which show similar modulatory effects on it. Note that this similarity is common, but not the same for all the subjects; for instance, ME-MOD showed larger reduction of motion related effects than ME-CON for two subjects (subject 003 and 007), while the opposite pattern was clearly observed in two other subjects (subject 004 and 009), and there was no apparent difference in the remaining subjects (see Supplementary figure 2). However, considering all subjects and all sessions together, the difference between these the ME-MOD and ME-CON approaches is statistically not significant (*β* =1.29, CI_95_ [-0.70, 3.27], p>0.5). Compared to OC-MPR, both ME-MOD and ME-CON reduce the impact of FD on DVARS (*β* =130.84, CI_95_ [127.89, 133.79], p<0.001 and *β* =132.13, CI_95_ [128.99, 135.27], p<0.001 respectively) The aggressive strategy (ME-AGG) is the most successful in reducing motion-related effects described by FD on DVARS of all approaches.

**Figure 3:**
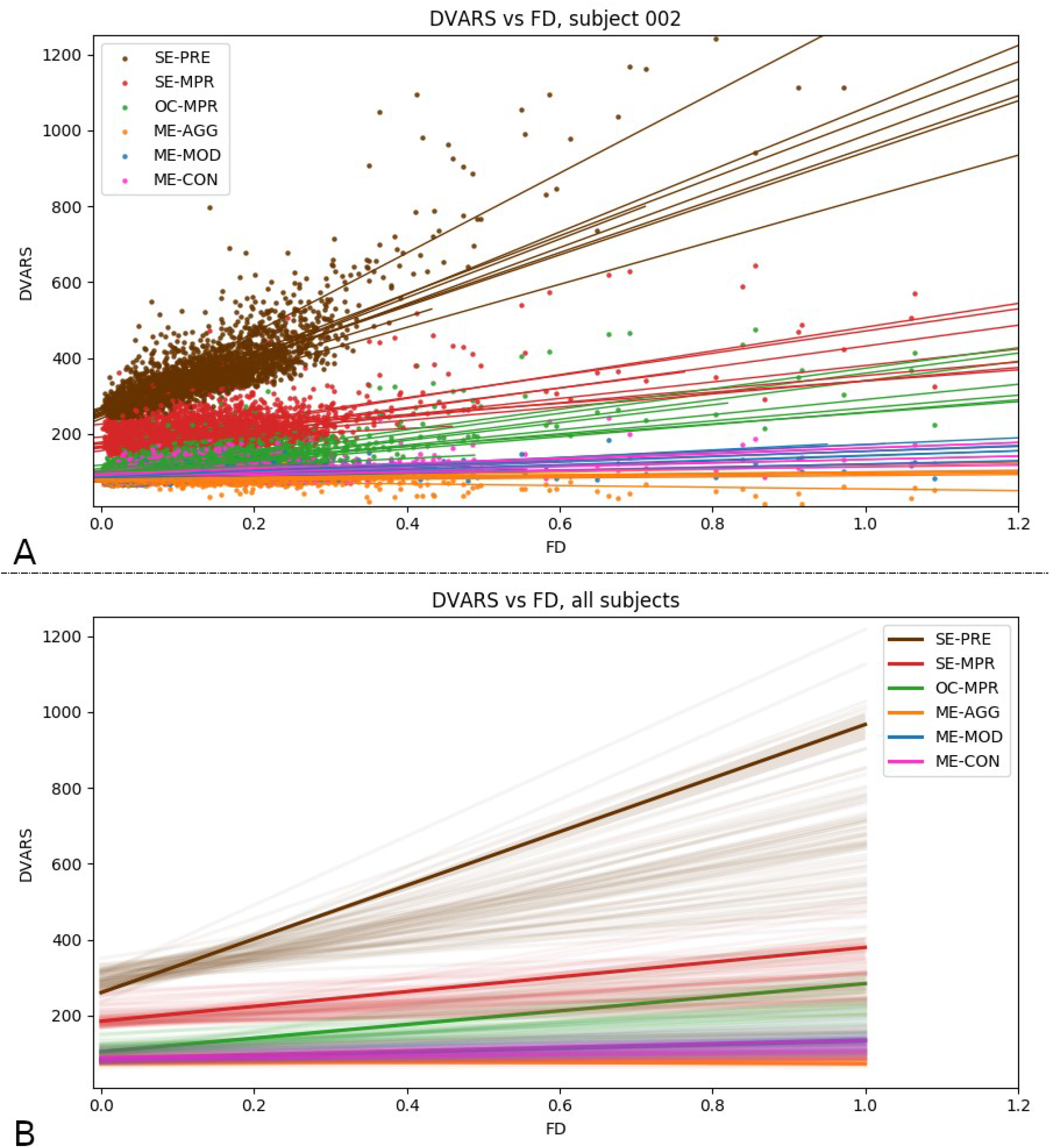
(A) Relation between the DVARS of the denoised data following different analysis pipelines and FD for a representative subject. Each point represents a timepoint, each line the linear regression between both timeseries in a session. In general, OC-MPR shows lower DVARS than SE-MPR, but similar modulation of the DVARS-FD relationship. All the ICA denoising solutions performs better in reducing motion-related effects described by FD on DVARS. Between the ICA solutions, ME-AGG performs the best in reducing this relationship, while ME-MOD and ME-CON seem to be equivalent. (B) DVARS vs. FD for all the subjects. Each transparent line represents a session, the solid line represents the estimation across subjects and sessions. Similar patterns to the representative subject are shown. SE-PRE: raw data; SE-MPR: single-echo; OC-MPR: optimally combined; ME-AGG: aggressive; ME-MOD: moderate; ME-CON: conservative. The relation between DVARS and FD of the other subjects can be found in the Supplementary Material (Supplementary figure 2).

**Table 1:** Comparisons of motion dependance in image intensity and general noise between different denoising approaches. * significant for p<0.001; all p values are computed with Satterhwaite’s method, and they are the equivalent of the p value after Šidák correction for multiple comparisons. SE-PRE: raw data; SE-MPR: single-echo; OC-MPR: optimally combined; ME-AGG: aggressive; ME-MOD: moderate; ME-CON: conservative.

|  | ME-AGG | ME-MOD | ME-CON | OC-MPR | SE-MPR |
| --- | --- | --- | --- | --- | --- |
| SE-PRE | $\beta = 715.10^*$,<br>CI<sub>95</sub> [710.17, 720.04] | $\beta = 658.05^*$,<br>CI<sub>95</sub> [653.09, 663.01] | $\beta = 659.34^*$,<br>CI<sub>95</sub> [654.27, 664.41] | $\beta = 527.21^*$,<br>CI<sub>95</sub> [521.74, 532.68] | $\beta = 512.65^*$,<br>CI<sub>95</sub> [506.88, 518.42] |
| SE-MPR | $\beta = 202.45^*$,<br>CI<sub>95</sub> [199.02, 205.88] | $\beta = 145.40^*$,<br>CI<sub>95</sub> [141.92, 148.88] | $\beta = 146.69^*$,<br>CI<sub>95</sub> [143.05, 150.33] | $\beta = 715.10^*$,<br>CI<sub>95</sub> [710.17, 720.04] | |
| OC-MPR | $\beta = 187.90^*$,<br>CI<sub>95</sub> [185.01, 190.78] | $\beta = 130.84^*$,<br>CI<sub>95</sub> [127.89, 133.79] | $\beta = 132.13^*$,<br>CI<sub>95</sub> [128.99, 135.27] | | |
| ME-CON | $\beta = 55.77^*$,<br>CI<sub>95</sub> [53.91, 57.63] | $\beta = 1.29$,<br>CI<sub>95</sub> [-0.70, 3.27] | | | |
| ME-MOD | $\beta = 57.05^*$,<br>CI<sub>95</sub> [55.59, 58.52] | | | | |

**Table 2:** Subject average CVR, lag, and percentage of statistical voxels in the grey matter across strategies. The last three lines are the group average. SE-PRE: raw data; SE-MPR: single-echo; OC-MPR: optimally combined; ME-AGG: aggressive; ME-MOD: moderate; ME-CON: conservative. The same table for the white matter is reported in the supplementary materials.

| Subject | Average value | SE-MPR | OC-MPR | ME-AGG | ME-MOD | ME-CON |
| --- | --- | --- | --- | --- | --- | --- |
| 001 | CVR [%BOLD/mmHg] | 0.54 | 0.5 | 0.37 | 0.43 | 0.49 |
|  | Lag [s] | -0.54 | -0.49 | 0.4 | -0.11 | -0.4 |
|  | % significant voxels | 9.79 | 10.44 | 3.33 | 8.1 | 10.75 |
| 002 | CVR [%BOLD/mmHg] | 0.38 | 0.35 | 0.24 | 0.3 | 0.35 |
|  | Lag [s] | -0.38 | -0.43 | 0.58 | 0 | -0.42 |
|  | % significant voxels | 10.67 | 11.65 | 3.29 | 8.61 | 11.92 |
| 003 | CVR [%BOLD/mmHg] | 0.4 | 0.34 | 0.18 | 0.31 | 0.33 |
|  | Lag [s] | -0.4 | -0.28 | -0.73 | -0.74 | -0.26 |
|  | % significant voxels | 7.42 | 8.04 | 3.62 | 6.55 | 8.28 |
| 004 | CVR [%BOLD/mmHg] | 0.44 | 0.38 | 0.08 | 0.32 | 0.37 |
|  | Lag [s] | -1 | -1.12 | -0.57 | -0.87 | -1.1 |
|  | % significant voxels | 8.46 | 9.27 | 2.87 | 6.57 | 9.56 |
| 007 | CVR [%BOLD/mmHg] | 0.33 | 0.29 | 0.17 | 0.28 | 0.29 |
|  | Lag [s] | -0.7 | -0.61 | 0.95 | -0.1 | -0.53 |
|  | % significant voxels | 7.98 | 9.19 | 2.6 | 6.31 | 9.44 |
| 008 | CVR [%BOLD/mmHg] | 0.34 | 0.14 | -0.03 | 0.26 | 0.14 |
|  | Lag [s] | -0.98 | -1.19 | -0.28 | -0.6 | -1.18 |
|  | % significant voxels | 6.34 | 6.89 | 1.5 | 4.9 | 7.19 |
| 009 | CVR [%BOLD/mmHg] | 0.44 | 0.38 | -0.18 | 0.31 | 0.37 |
|  | Lag [s] | -1.75 | -1.75 | 0.91 | -0.11 | -1.69 |
|  | % significant voxels | 7.52 | 9.16 | 2.25 | 6.42 | 9.55 |
| Total | CVR [%BOLD/mmHg] | 0.41 | 0.34 | 0.12 | 0.32 | 0.33 |
|  | Lag [s] | -0.82 | -0.84 | 0.18 | -0.36 | -0.80 |
|  | % significant voxels | 8.31 | 9.23 | 2.78 | 6.78 | 9.53 |

Figure 4a plots the average percentage DVARS (left column) and average GM percentage BOLD response (central column) of all the BH trials across all of the sessions of a representative subject. The FD trace features a clear peak right after the end of the apnoea (highlighted in grey), likely associated with large head movement artefacts caused by the recovery breaths following the apnoea period. The percentage DVARS curves of the SE-PRE, SE-MPR and OC-MPR denoised timeseries reflect this peak in FD, which is absent in the ME-ICA based denoising timeseries, indicating a strong influence of movement on the signal intensity changes. All DVARS curves present a peak at a later time (between timepoints 25 and 30) that, as DVARS is akin to the first derivative of the BOLD signal changes, may agree with the return to the baseline seen in the BOLD response. The percentage BOLD signal change curves feature a delayed peak compared to the FD trace, reflecting a delayed CVR response compared to instantaneous head movements associated with respiration. However, they also feature a modulation in the BOLD signal change in correspondence with the peak in the FD trace, with the exception of ME-MOD and ME-AGG. The flattened DVARS and BOLD responses seen for ME-AGG indicate that the inclusion of the ME-ICA rejected components substantially removes part of the true CVR response, compared with the OC-MPR time courses. The average percentage DVARS and percentage BOLD response of the other subjects can be found in the Supplementary Materials (Supplementary figure 3).

**Figure 4:**
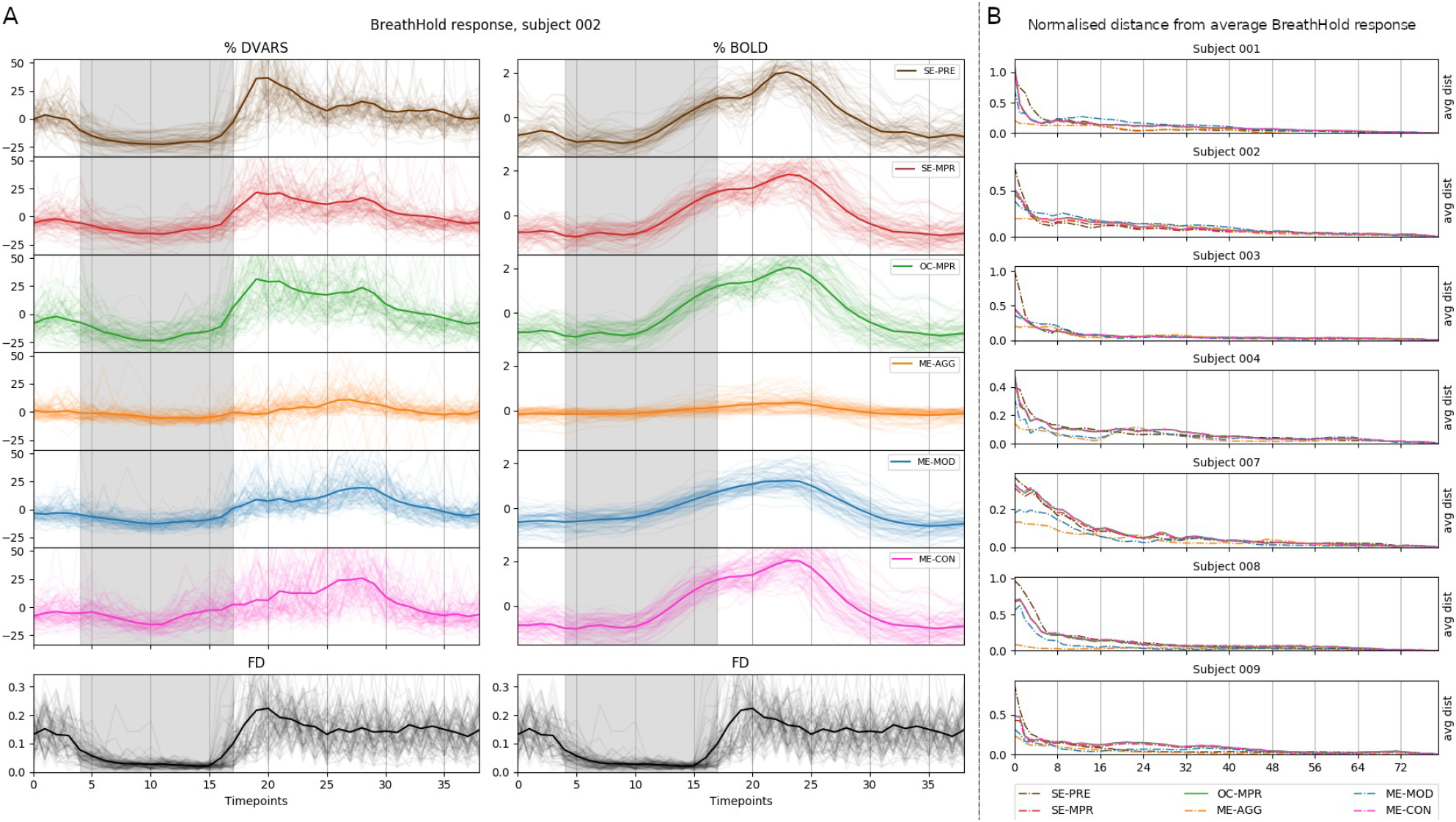
(A) Average GM %DVARS and %BOLD response of all BH trials across ten sessions for the same representative subject. The apnoea period is highlighted in grey. Each transparent line is a trial, the solid line is the average across all the trials. (B) Manhattan distance between the average of N trials and the average of all 80 BH trials as N increases from 1 to 80 for each subject. Each vertical line divides the number of trials in each session. SE-PRE: raw data; SE-MPR: single-echo; OC-MPR: optimally combined; ME-AGG: aggressive; ME-MOD: moderate; ME-CON: conservative. The average %DVARS and %BOLD response of the other subjects can be found in the Supplementary Material (Supplementary figure 2)

Figure 4b plots the Manhattan distance between the average of *N* trials and the average of all 80 BH trials as *N* increases from 1 to 80. ME-AGG tends to be more similar to the total average compared to all the other timeseries. For most of the subjects, SE-MPR, OC-MPR and ME-MOD have a similar behaviour and need more trials than SE-PRE, ME-CON and ME-AGG to converge to the total average. Note that the convergence to the analysis-specific ‘ground truth’ BH response is not monotonic and fluctuates across trials of the same session and across sessions, indicating that the convergence does not depend only on the number of BH trials, but also on their quality and possible physiological variability in the CVR response across trials and sessions.

### 3.2 Cerebrovascular reactivity and lag maps

Figure 5 and 6 show CVR and lag maps respectively, for all analysis strategies and all sessions of a representative subject (subject 002). The CVR and lag maps of other subjects are available in the *S*upplementary Material (Supplementary figure 4 and 5). The CVR maps were masked to exclude the voxels that were not statistically significant or whose lag is at the boundary of the explored range and might not have been truly optimised or physiologically plausible. Across all subjects, SE-MPR features more spatial variation and speckled noise in CVR and lag estimates of voxels within the same brain region compared to ME approaches like OC-MPR or ME-CON. In general, the ME-AGG and ME-MOD approaches do not yield CVR maps with as much clear distinction between brain tissues or delineation of the cortical folding and subcortical structures (e.g. see putamen and caudate nucleus) as obtained with the OC-MPR and ME-CON models. Among the ICA-based approaches, the adoption of an aggressive (ME-AGG) or moderate (ME-MOD) modelling strategy results in lag maps without anatomically defined patterns, as well as a higher rate of voxels with a lag estimation that is not within physiologically plausible range, and in CVR maps with lower responses and fewer significant voxels. ME-AGG also produces CVR maps with a higher percentage of negative values than any other analysis model, and a reduced CVR response in voxels near the posterior part of the superior sagittal and transverse sinuses.

**Figure 5:**
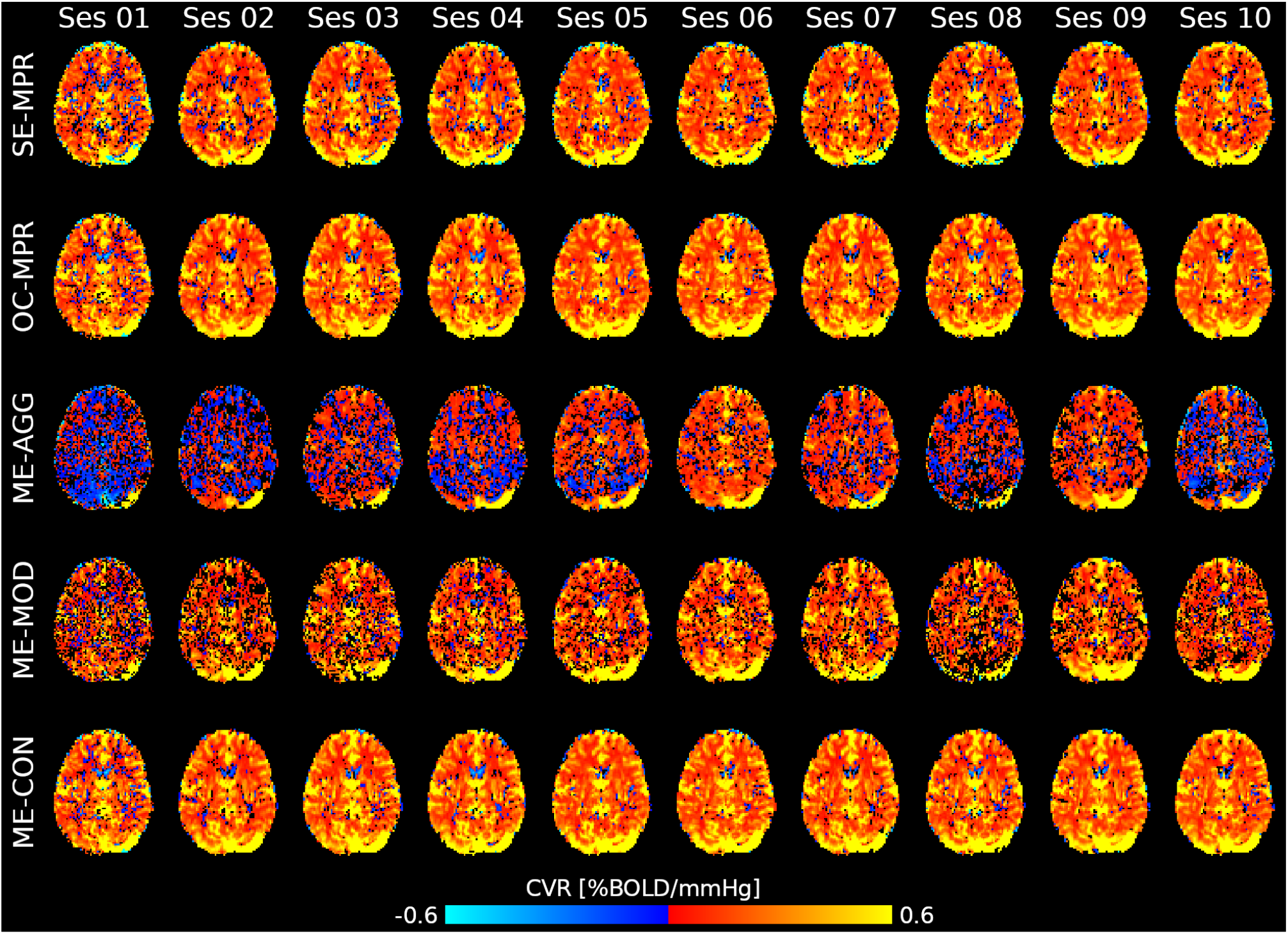
Thresholded CVR map obtained with the different lagged-GLM analysis for all the sessions of a representative subject (subject 002). Note the low CVR response in ME-AGG, depicting numerous voxels with a negative values, as well as the increased amount of masked voxels in SE-MPR, ME-AGG and ME-MOD. SE-MPR: single-echo; OC-MPR: optimally combined; ME-AGG: aggressive; ME-MOD: moderate; ME-CON: conservative. The CVR maps of other subjects are available in the supplementary materials.

**Figure 6:**
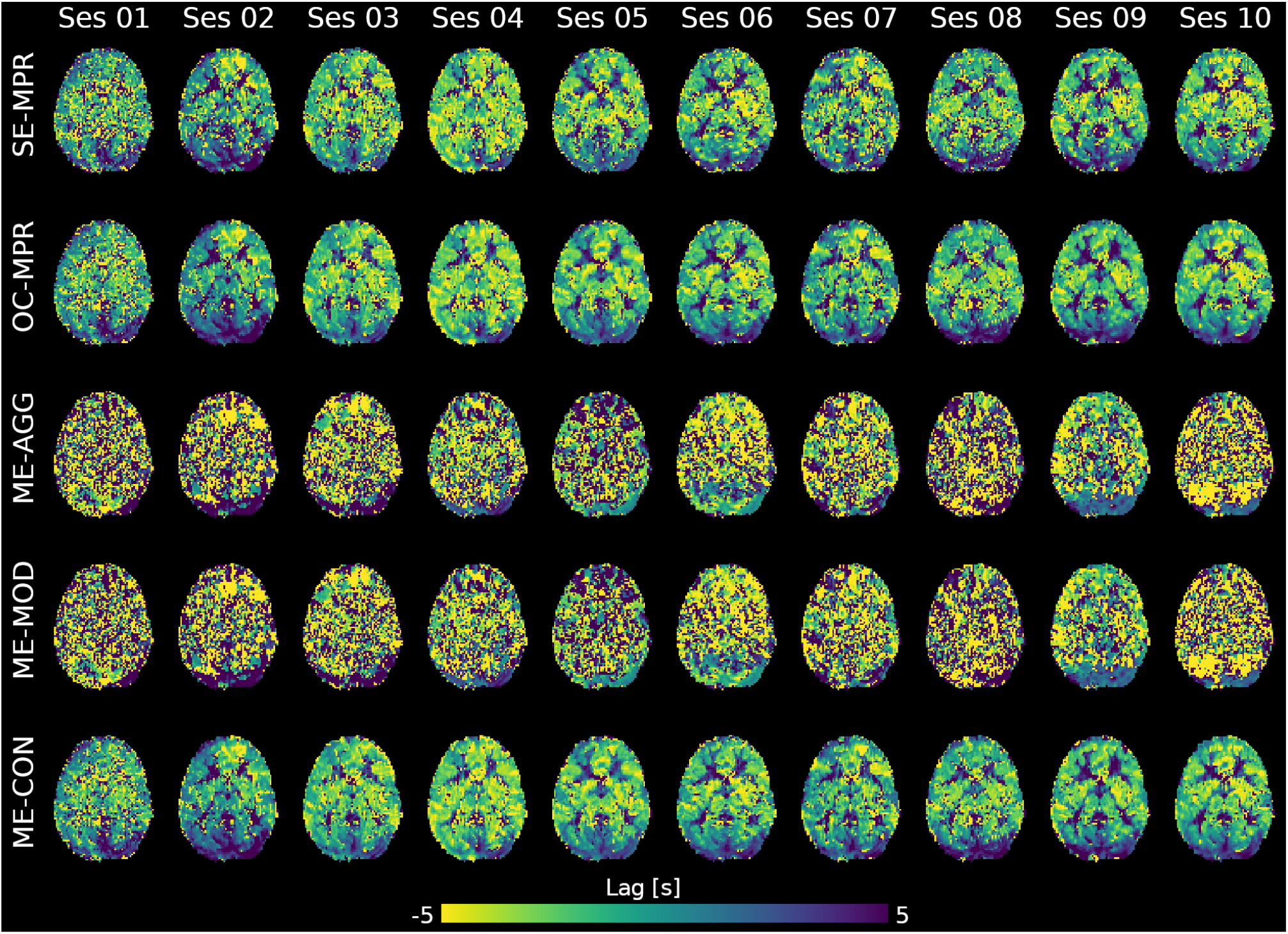
*Unthresholded lag map obtained with the different lagged-GLM analysis, for all the sessions of a representative subject (same as figure 5). These lag maps represent the delay between the best shifted version of the* P_ET_CO_2_*hrf trace and the bulk shift (i.e. the best match between average grey matter signal and* P_ET_CO_2_*hrf trace). The scale from −5 to +5 represents earlier to later hemodynamic responses. Note the lack of anatomically informative patterns in ME-MOD and ME-AGG. SE-MPR: single-echo; OC-MPR: optimally combined; ME-AGG: aggressive; ME-MOD: moderate; ME-CON: conservative. The lag maps of other subjects are available in the supplementary materials*.

Figure 7 shows the distribution of the average values of CVR, lag, and the percentage of significant voxels for all subjects and sessions, and across all denoising strategies after thresholding. Considering the summaries within GM, although SE-MPR shows higher average CVR compared to the other approaches, it also features lower percentage of significant voxels compared to OC-MPR, ME-MOD and ME-CON. ME-AGG shows the lowest CVR value of all strategies, the most variable average of lag values, as well as the lowest percentage of significant voxels. ME-MOD features a lower percentage of significant voxels than SE-MPR, OC-MPR, and ME-CON. The same considerations can be extended to the WM. Table 2 reports the subject average CVR, lag, and the percentage of significant voxels across all denoising strategies after thresholding for GM only. The same table for WM can be found in the supplementary materials (Supplementary table 1). For all models, the average CVR in the GM in the group and in each subjects are comparable or higher than the reported BH-induced CVR (in %BOLD/mmHg) in previous literature (cfr. Bright et al., 2011; Bright & Murphy, 2013a; Lipp et al., 2015; Pinto et al., 2016).

**Figure 7:**
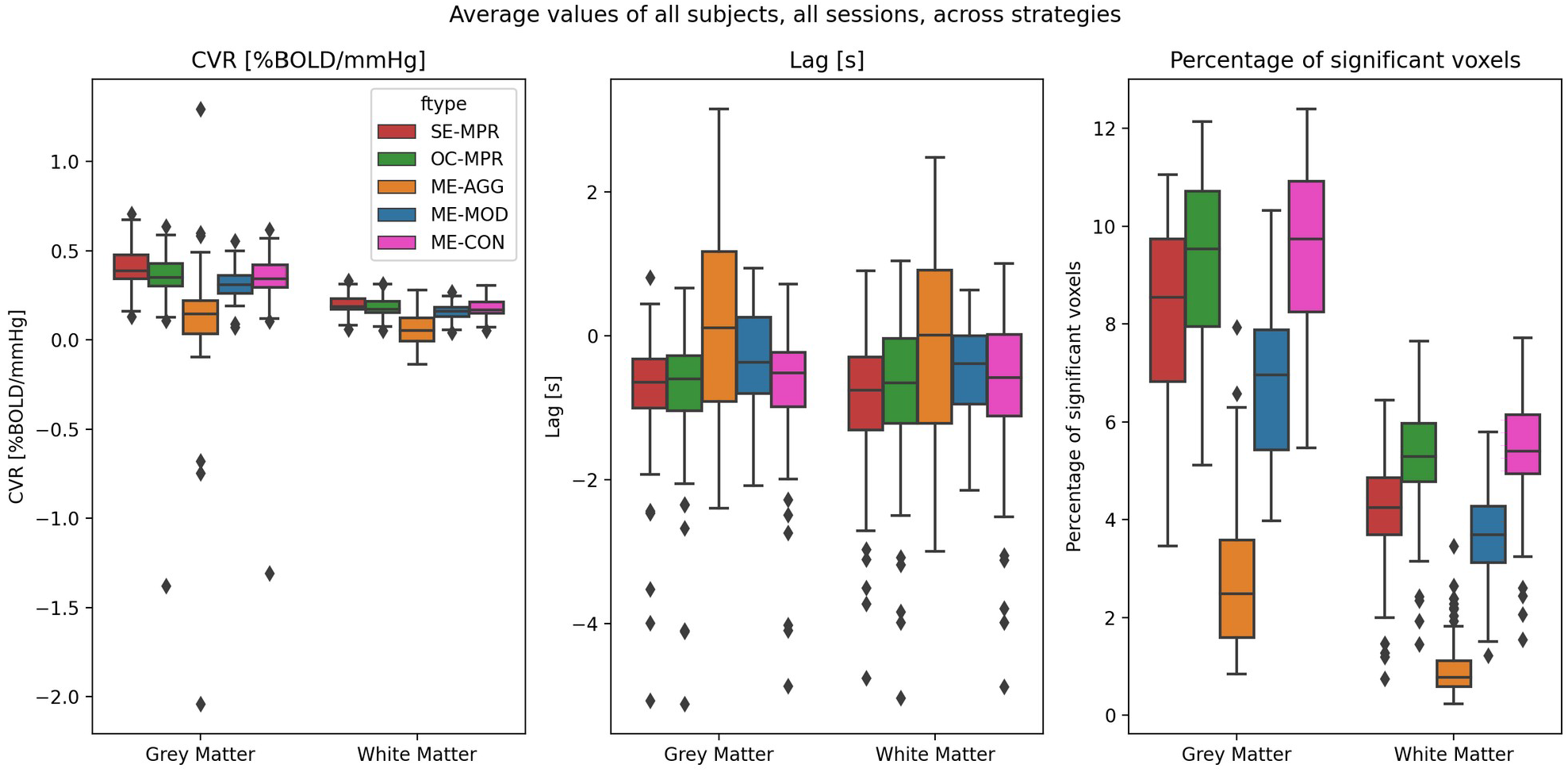
Average values of CVR, lag, and percentage of significant voxels, for voxels in the grey and white matter tissues separately, for all denoising strategies. The dots correspond to a singular session of a singular subject considered an outlier in the distribution. Note that all maps were thresholded before plotting. SE-MPR: single-echo; OC-MPR: optimally combined; ME-AGG: aggressive; ME-MOD: moderate; ME-CON: conservative.

### 3.3 Comparison of CVR and lag estimation and reliability across denoising strategies

Figure 8 shows the results of comparing the CVR and lag maps across all of the denoising strategies. The top row shows the thresholded *χ* score of the contrast between SE-MPR and all other denoising strategies, while the other maps depict the pairwise comparison between all of the denoising strategies. Among the most interesting comparisons, all of the strategies based on ME have lower CVR and an anticipated response in areas vascularised by big vessels (indicated by an arrow in the figure), where the blood transit time is usually faster compared to the rest of the brain. This could indicate that the response shown in SE-MPR could be overestimated due to the misestimation of its lag. Compared to SE-MPR, ME-MOD shows lower CVR and a delayed response in subcortical areas, while OC-MPR and ME-CON show higher CVR and an anticipated response in the insula, frontal, and parietal areas. OC-MPR shows no statistically significant differences with ME-CON, but a general higher CVR and an anticipated response compared to ME-MOD and ME-AGG, with the exception of the cerebellum, where it shows a delayed response. This difference could be related to the different local impact of motion artefacts, especially on the cerebellum. Between the three approaches based on ME-ICA, ME-AGG features generally lower CVR compared to the other two, and a generally anticipated response compared to ME-MOD and a delayed response to ME-CON.

**Figure 8:**
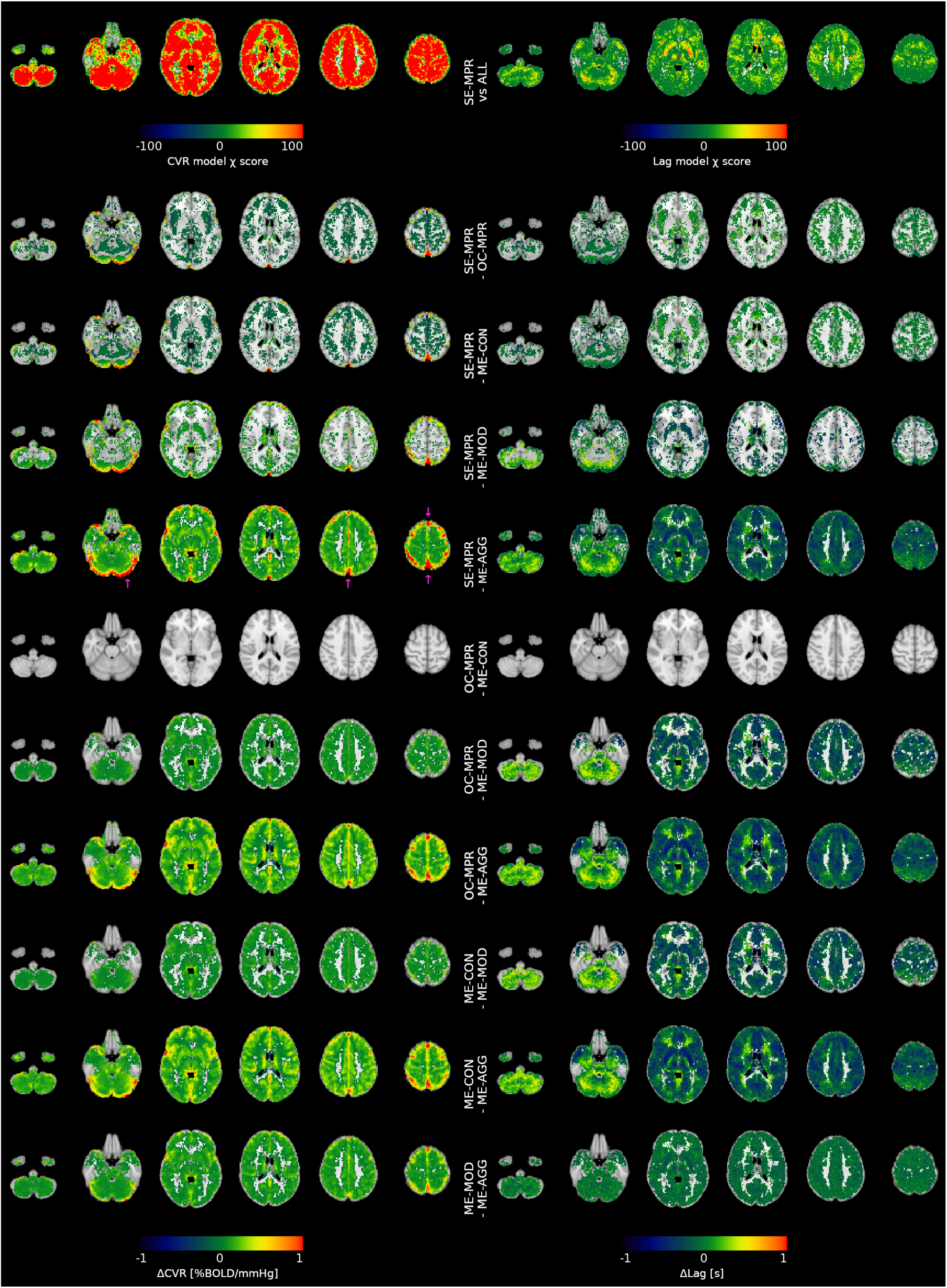
Top row: Thresholded χ value of the LME model used for the comparison of CVR and lag maps across all denoising strategies. Arrows indicate areas vascularised by big vessels. Other rows: pairwise comparison between denoising strategies. Arrows indicate SE-MPR: single-echo; OC-MPR: optimally combined; ME-AGG: aggressive; ME-MOD: moderate; ME-CON: conservative.

In order to assess the reliability of each model, we also computed voxelwise ICC(2,1) maps for both CVR and haemodynamic lag. Figure 9 depicts the ICC(2,1) maps for all analysis strategies for both CVR and lag maps, as well as their distributions. High ICC scores indicate that the intra-subject variability is lower than the inter-subject variability, hence the estimations of CVR or haemodynamic lag can be considered consistent across sessions. Conversely, low ICC scores indicate that the inter-subject variability is low compared to the intra-subject variability, hence the estimations of CVR and haemodynamic lag cannot be considered consistent across sessions. Following the classification given by (Cicchetti, 2001), an ICC score lower than 0.4 is considered poor, lower than 0.6 fair, lower than 0.75 good, and equal or higher than 0.75 excellent.

**Figure 9:**
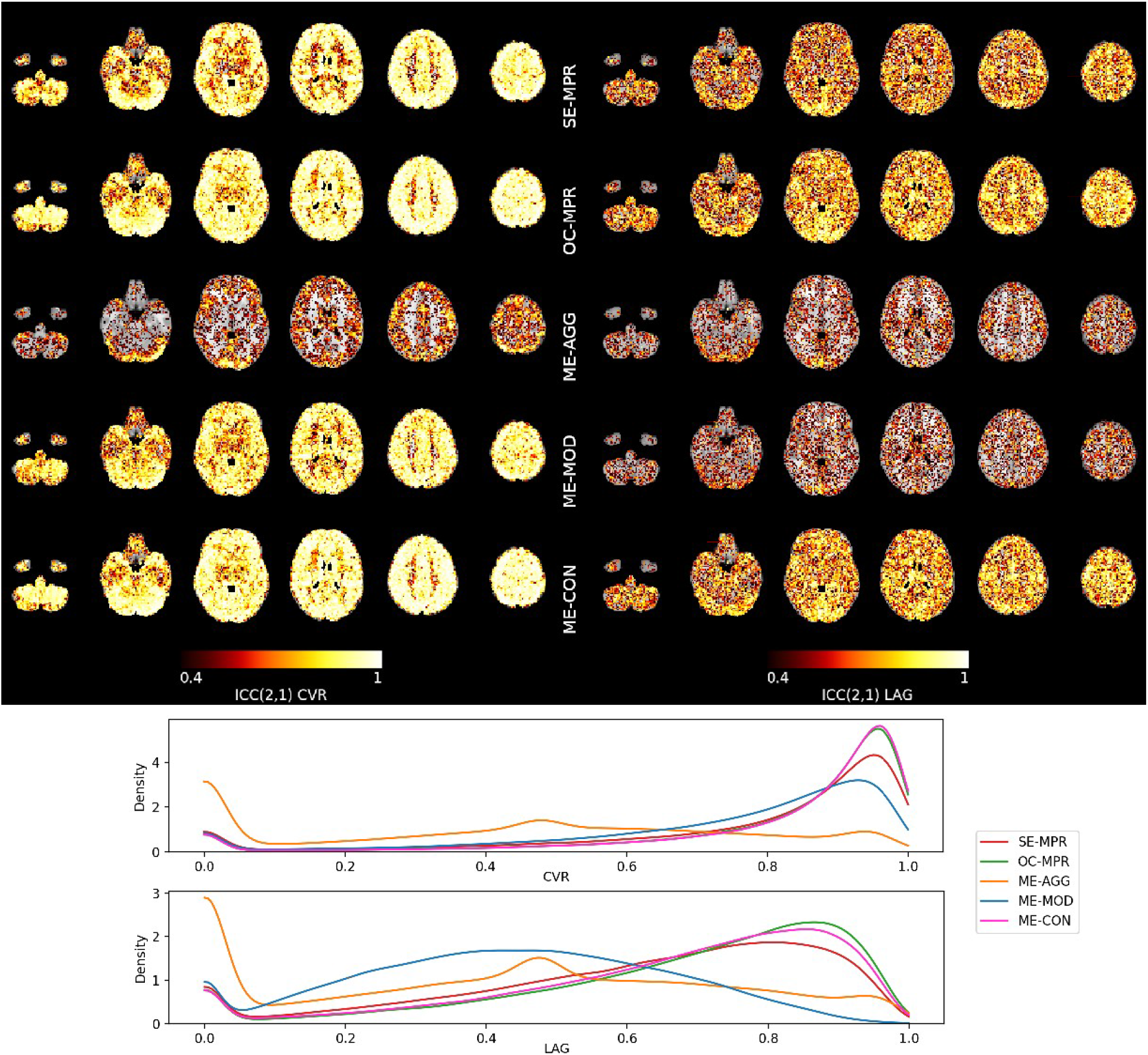
ICC(2,1) maps of CVR (left) and haemodynamic lag (right) for each analysis pipeline. The maps are thresholded at 0.4 since scores lower than it indicate poor reliability. A high ICC score indicates that the inter-subject variability is higher than the intra-session variability, while a low ICC score suggest that the variability across sessions is the same as the one across subjects. Following the classification given by Cicchetti (2001), an ICC score lower than 0.4 is considered poor, lower than 0.6 fair, lower than 0.75 good, and equal or higher than 0.75 excellent. The bottom rows depict the whole brain distribution of ICC scores across voxels. Note how OC-MPR and ME-CON have generally higher ICC scores than the other approaches, and are very similar to each other, while ME-AGG has the lowest ICC scores for both CVR and lag maps. SE-MPR: single-echo; OC-MPR: optimally combined; ME-AGG: aggressive; ME-MOD: moderate; ME-CON: conservative. The distribution of ICC scores across GM voxels only is available in the Supplementary Material (Supplementary figure 6).

In terms of whole brain CVR reliability, the ME-CON demonstrated excellent reliability (spatial average across the whole brain of 0.86 ± 0.16) as well as the highest ICC values among all methods tested, closely followed by the OC-MPR (excellent, 0.85 ± 0.16), SE-MPR (excellent, 0.81 ± 0.19), and ME-MOD (excellent, 0.79 ± 0.19), while ME-AGG had a fair reliability (0.46 ± 0.22). If only voxels in GM are considered, the ICC of all approaches increases slightly (0.88 ± 0.14, 0.87 ± 0.15, 0.85 ± 0.17, 0.82 ± 0.17, and 0.49 ± 0.22 for ME-CON, OC-MPR, SE-MPR, ME-MOD, and ME-AGG respectively). Despite the average fair reliability observed for ME-AGG, it can be observed that this approach exhibits a considerable number of voxels with poor reliability (ICC below 0.4). These voxels are mostly located in white matter, which also exhibit lower ICC values in the other analyses. In terms of whole-brain lag reliability, OC-MPR performed the best (good reliability, 0.67 ± 0.21), closely followed by ME-CON (good reliability, 0.66 ± 0.21). SE-MPR, ME-MOD, and ME-AGG demonstrated fair lag reliability (0.6 ± 0.22 and 0.42 ± 0.19, 0.41 ± 0.20, respectively). Considering only GM voxels, the reliability of all the approaches increases minimally (0.68 ± 0.21, 0.67 ± 0.21, 0.61 ± 0.21, 0.43 ± 0.19, 0.42 ± 0.20, for OC-MPR, ME-CON, SE-MPR, ME-MOD, and ME-AGG respectively). The reliability of CVR lag estimates was lower than that of CVR amplitude estimates, even though certain cortical regions, such as the visual and motor cortices, also show excellent ICC values for the OC-MPR and ME-CON denoising approaches. Interestingly, it can be observed that ME-MOD offers excellent ICC values for the CVR response amplitude in grey matter voxels, whereas they are poor for the lag estimates.

## 4 Discussion

In this study, we compared five different analysis strategies based on a lagged-GLM model (Moia, Stickland, et al., 2020) to simultaneously remove motion-related effects and non-BOLD artefacts in the BOLD fMRI signal while estimating CVR and haemodynamic lag in order to identify the best modelling approach for BH paradigms in which prominent task-correlated artefacts coexist with the effect of interest. The lagged-GLM model adopted in this study is similar to other models for CVR estimation that take into account local variations in the haemodynamic lag (Donahue et al., 2016; Geranmayeh et al., 2015; Murphy et al., 2011; Sousa et al., 2014; Tong et al., 2011). Since a comparison with the other models was outside of the scope of this study, we did not perform it. Further studies should perform such comparisons and observe how an ICA denoising approach should be adapted to them.

Among all possible modelling strategies, the five presented here were included in our analysis for different reasons. The optimal combination of ME fMRI data, with subsequent motion and Legendre polynomial regression (MPR), was expected to remove more noise and improve reliability of the CVR estimation due to its increased BOLD sensitivity compared to MPR on single-echo data, which is the standard approach for BH CVR estimation (Cohen & Wang, 2019). However, while optimal combination of ME volumes alone can partially reduce the noise present in the data, it still cannot remove motion artefacts, as illustrated in Figure 3 in which SE-MPR and OC-MPR exhibit the same dependence of signal changes (DVARS) with motion (FD). For this reason, we further adopted three different ME-ICA based approaches, ranging from a conservative to an aggressive motion removal. ICA-based approaches are known to outperform traditional MPR in typical denoising fMRI data, possibly because they can identify and separate artefactual sources in the data in a data-driven and non-linear manner (Griffanti et al., 2014; R. H. R. R. Pruim et al., 2015; Salimi-Khorshidi et al., 2014). We did not apply ICA to single-echo data because it has already been demonstrated that ICA-based denoising applied to OC data outperforms ICA denoising applied to single-echo data (Dipasquale et al., 2017) and the ICs estimated from OC data might not have matched the ICs obtained from single-echo data, making such comparison less straight forward than the one based on MPR.

Spatial ICA decomposition is applied to fMRI data more often than temporal ICA decomposition, as the latter requires many more samples in time than normally available. Having many sessions for each subject, temporal ICA could have been leveraged in this study. In fact, temporal ICA could be more appropriate than spatial ICA to estimate a proper decomposition of timeseries sources (Smith et al., 2012), improving the modelling of temporal noise (Glasser et al., 2018), and potentially leading to better disentanglement of noise from CVR effects. However, we decided to apply spatial ICA in order to maintain the independence of each session, both to simulate a more common denoising approach to fMRI data, and to be able to capture session-specific noise contributions that could have been missed otherwise. Further studies could compare temporal and spatial ICA denoising for CVR mapping when many temporal samples have been collected in the same session, for instance reducing the TR by acquiring fewer echoes. Here, our decision to acquire five echoes, instead of conventional multi-echo protocols with three or four echoes, was made to facilitate and improve the classification of the ICs based on their TE-dependence (Kundu et al., 2013).

The choice of comparing different levels of orthogonalisation of only the ICA-based nuisance regressors compared to regressors of interest might seem in contrast with previous literature, that suggests that orthogonalisation of collinear confounding factors could lead to misinterpreted results (Mumford et al., 2015). Our results clearly demonstrated that using the original (e.g., non orthogonalised) rejected ICs as nuisance regressors in the analysis (ME-AGG) removes the CVR effect of interest (see Figures 4, 5, 6 and 9). To decide which regressors should be orthogonalised, and with respect to what, we considered the different origin of the nuisance regressors. While Legendre polynomials and motion parameters can be considered adequate models of noise sources in the data, intrinsic data-driven regressors may well contain variance related to the effect of interest, especially as spatial ICA was adopted and because of the high collinearity between the P_ET_CO_2_*hrf*, motion, physiological adaptations to vascular dilation (e.g. cerebrospinal fluid flows), or changes in the magnetisation related to breathing (Raj et al., 2001). In these scenarios, it becomes more important to understand how to properly implement ICA denoising in order to preserve the effect of interest. For these reasons, three different ICA-based approaches were selected, from an aggressive strategy to a conservative approach, to assess if they preserved the BOLD effects related to the CVR response happening at different lags.

As hypothesized, all of the ME-based solutions outperformed the SE-MPR model in their ability to account for the effect of motion, summarized in terms of FD, on the fMRI signal intensity changes, described in terms of DVARS (see Figure 3). Furthermore, all of the ICA-based strategies outperformed traditional MPR, and within ICA-based strategies, the aggressive one (ME-AGG) showed the best performance to remove these motion-related effects in the signal. However, observing the average DVARS and BOLD response timecourses (Figure 4) and the CVR and lag maps (Figures 5 and 6) it becomes evident how aggressive and moderate approaches result in lower estimates of CVR responses, even compared to the SE-MPR approach. Similarly, these two approaches result in the estimated haemodynamic lag hitting the boundaries of a physiologically plausible lag range in healthy adults. The substantial reduction in the CVR estimates in the aggressive approach (Figures 4 and 5) occurs because the effect of interest can also be explained as a linear combination of the timecourses of rejected ICs related to motion, vascular effects or large susceptibility changes due to chest expansions and contractions while performing the BH task (Caballero-Gaudes & Reynolds, 2017; Griffanti et al., 2017). As for the moderate approach, the lower estimates of CVR could be due to the fact that orthogonalising data-driven nuisance regressors with respect to the P_ET_CO_2_*hrf* trace per sé is not sufficient to save all the variance associated to real CVR. The P_ET_CO_2_ trace can only be estimated during exhalations, hence it is unable to capture local dynamic signal changes that are captured by ICs timeseries. Furthermore, CVR has a sigmoidal non-linear relation with the P_ET_CO_2_*hrf* trace (Bhogal et al., 2014), and the local BH-induced BOLD response has a complex shape, in terms of response amplitude and temporal delays, due to multiple physiological factors (Magon et al., 2009) that must be accounted for in order to improve its estimation. Our results illustrate that these local complexities might be adequately captured by the linear combination of the accepted ICs timecourses, and not removing this variance from the rejected ICs when they are included as nuisance regressors in the model is detrimental (as observed with ME-MOD and ME-AGG approaches). In other words, only a conservative approach (ME-CON) that preserves the BOLD variance associated with local CVR responses performs well, while also reducing motion-related effects more than conventional MPR models.

To further explore the benefit of different modelling strategies, we assessed the reliability of the resulting CVR and haemodynamic lag maps over the course of two and a half months (ten sessions) using ICC(2,1). To our knowledge, this was the first time that CVR reliability was tested over the course of ten sessions in individual subjects, and the first time that intersession haemodynamic lag reliability was tested. The ME-CON and OC-MPR strategies featured the greatest reliability for CVR and lag estimation, while the ME-AGG and ME-MOD approaches produced lower reliability values than even the simple SE-MPR model.

The lag maps are computed as the temporal offset related to the bulk shift, which is obtained by aligning the average GM BOLD response with the P_ET_CO_2_*hrf* trace. If the bulk shift computation is misestimated this would create a systematic bias in the estimated lag maps, potentially reducing the apparent intersession reliability. While the CVR reliability should not be affected by this issue, due to the use of a lagged-GLM approach that can overcome bulk shift misestimation (see session 4 of subject 007 in Supplementary figure 4 and 5), the true lag map reliability might be higher than reported here.

Regarding CVR reliability, the whole-brain average reliability of SE-MPR was comparable to long-term reliability (days or weeks apart) found in previous studies of CVR induced by BH (Peng et al., 2019), by paced deep breathing (Sousa et al., 2014), or by gas challenges (Leung et al., 2016), and higher than that reported in other studies on BH induced CVR estimated with a non-lagged optimized P_ET_CO_2_*hrf* trace (Lipp et al., 2015) or with Fourier modelling (Pinto et al., 2016), and by gas challenges (Dengel et al., 2017; Evanoff et al., 2020). Consequently, the reliability of CVR estimates obtained with the optimal combination dataset and conservative ME-ICA modelling approaches were found higher than those previously reported in the literature. However, all strategies produced a reliability that was lower than the short*-*term (within-session) reliability reported in BH induced CVR (Peng et al., 2019), resting state based CVR (P. Liu et al., 2017), and gas challenge induced CVR (Leung et al., 2016), although lower intersession reliability in gas challenges has also been reported (Dengel et al., 2017; Evanoff et al., 2020). Note that the reliability observed in this study seems to be globally higher and spatially less variable than that reported in previous studies (Lipp et al., 2015; Sousa et al., 2014). However, discrepancies in the reliability measurements might be related to the different methods used to compute the CVR maps and the ICC score itself.

Using ICC to test reliability has the drawback that higher scores might be related to the presence of residual task-correlated motion effects that artificially stabilise the CVR estimation and reduce intrasubject variability compared to intersubject variability. In fact, recent studies have shown that individuals have particular movement patterns during fMRI sessions that may be a stable characteristic of a person (Bolton et al., 2020) related to stable physical characteristics, such as body mass index (Ekhtiari, Kuplicki, Yeh, & Paulus, 2019) and could even be a heritable characteristic (Couvy-Duchesne et al., 2014; Hodgson et al., 2017). If subjects have similar motion patterns across the 10 repeated sessions, fMRI responses might appear more similar than they truly are, and the ICC might be inflated by such effects. Moreover, higher spatial reliability does not necessarily mean higher accuracy: a denoising strategy might be systematically misestimating CVR or haemodynamic lag. The fact that both optimal combination with traditional nuisance regression and the conservative ME-ICA denoising approaches resulted in similar CVR and lag spatial patterns and exhibited higher reliability than the single-echo model, while at the same time reduced the apparent effect of motion on the data variance, suggests that such drawbacks are mitigated in our data. However, further studies could compare different BH analysis strategies with a CVR estimation based on an independent computerised gas delivery protocol.

Another possibility would be to assess CVR in resting state fMRI, either measuring resting fluctuations in exhaled CO_2_ levels (Golestani, Wei, & Chen, 2016; Lipp et al., 2015), or by using a band of the power spectrum of the global signal as a regressor of interest (P. Liu et al., 2017, 2020). Such method might be more robust to motion collinearity, as the amount of movement in each breath is less pronounced and not consistently time-locked to the paradigm cues. At the same time, the lower amplitude of intrinsic CO_2_ fluctuations relative to BH CO_2_ change might also make this approach more susceptible to general motion effects and other sources of variance (e.g. neural or artefactual) unrelated to CO_2_. Moreover, previous work has shown that the optimal temporal shift between BOLD and P_ET_CO_2_ is hard to reliably identify in resting state data alone, in contrast to BH datasets where the temporal shift can be reliably identified (Bright et al., 2017; Stickland, Ayyagari, Zvolanek, & Bright, 2020). Current resting state fMRI methods for CVR mapping may therefore be inappropriate to use with the lagged-GLM approach that we have applied here. Either way, the analyses presented in this study can be easily implemented in other CVR assessment pipelines to mitigate the dependence of the response on motion. Beyond BH-based CVR studies, similar conclusions might be applicable to other experimental paradigms that present high collinearity between the expected task induced activity and artefactual sources, such as in overt speech production with long trial durations (Birn et al., 1999, 2004; Gracco, Tremblay, & Pike, 2005), that aim to use (ME-) ICA-based nuisance regressors as part of the model.

Note that MPR and ICA denoising are not the only viable options to reduce motion effects on fMRI and BH-induced CVR in particular: advanced setups can be used to reduce motion during the acquisition itself. For instance, subject specific moulded head casts can be used to reduce head motion (Power, Silver, et al., 2019). Mounting an MRI compatible camera or tracker in the scanner enables prospective motion correction techniques (Faraji-Dana, Tam, Chen, & Graham, 2016; Maziero, Rondinoni, Marins, Stenger, & Ernst, 2020; Parkes, Fulcher, Yücel, & Fornito, 2018; Schulz et al., 2014; Zaitsev, Akin, LeVan, & Knowles, 2017) or concurrent field monitoring enables the dynamic correction of field distortions dynamically (Vannesjo et al., 2015; Wilm et al., 2015) in order to effectively reduce effects of motion and magnetic field susceptibility changes. However, such advanced setups are not always available.

A limitation of this study is that the results are influenced by the manual classification of the ICA components performed by two of the authors. Despite being based on the automatic classification made by *tedana*, we adopted a manual approach because often multiple ICs clearly exhibiting CVR-related timeseries and spatial maps were misclassified as noise. This manual classification was made with a cautious approach: if an IC seemed to be temporally and spatially related to the CVR response, it was accepted. Manual classification is still considered the gold standard for the classification of ICA components when performed by experts, despite the introduction of automatic classification algorithms (Griffanti et al., 2017), calling for further improvements in the automatic classification of (ME-)ICA components for BH tasks.

Moreover, despite the fact that a BH task can be a valid alternative to gas delivery protocols for CVR estimation and its easy implementation, not all the subjects in this study could perform the task during all of the sessions. In total, 86% of the sessions were completed successfully by the subjects, although three subjects had to be excluded due to poor performance or non-compliance to the task in a subset of the sessions (four in two subjects and six in the third, see Figure 2).

Finally, it is worth noticing that the adoption of ME imaging requires an increase in TR or a decrease in the spatial resolution. A way to cope for this issue is the adoption of simultaneous multislice (a.k.a. multiband) acquisition, and despite the fact that this choice might introduce additional slice-leaking artefacts, a ME-ICA based denoising approach can successfully deal with their removal (Olafsson, Kundu, Wong, Bandettini, & Liu, 2015). Note that in this study we adopted one of the echo volumes as an approximation of a SE acquisition. Further studies could evaluate if this solution improves the estimation of CVR compared to SE imaging with higher spatial or temporal resolution.

## 5 Conclusion

Breath Holding (BH) is a non-invasive, robust way to estimate cerebrovascular reactivity (CVR). However, due to the task-correlated movement introduced by the BH task, attention has to be paid when choosing an appropriate modelling strategy to remove movement-related effects while preserving the effect of interest (P_ET_CO_2_). We compared different multi-echo (ME) independent component analysis (ICA) based denoising strategies to the standard data acquisition and analysis procedure, i.e. single-echo motion parameters regression. We found that a conservative ICA-based approach, but not an aggressive or moderate ICA approach, best removes motion-related effects while obtaining reliable CVR and lag responses, although a simple optimal combination of ME data with motion parameters regression provides similar CVR and lag estimations, and both ME-based approaches offer improvements in reliability compared with single-echo data acquisition.

## 6 Acknowledgements

CRediT statement. Stefano Moia: Conceptualisation, Methodology, Software, Formal Analysis, Investigation, Data Curation, Writing (OD), Writing (RE), Visualisation, Funding acquisition; Maite Termenon: Methodology, Supervision, Writing (RE); Eneko Uruñuela: Investigation, Writing (RE); Rachael C. Stickland: Methodology, Writing (RE); Gang Chen: Methodology, Formal Analysis, Writing (RE); Molly G. Bright: Methodology, Supervision, Resources, Writing (RE); César Caballero-Gaudes: Conceptualisation, Methodology, Investigation, Supervision, Resources, Writing (RE), Project Administration, Funding acquisition.

The authors would like to thank Vicente Ferrer for collaborating in data acquisition and two anonymous reviewers for helping improving the quality of the paper.

This research was supported by the European Union’s Horizon 2020 research and innovation program (Marie Skłodowska-Curie grant agreement No. 713673), a fellowship from La Caixa Foundation (ID 100010434, fellowship code LCF/BQ/IN17/11620063), the Spanish Ministry of Economy and Competitiveness (Ramon y Cajal Fellowship, RYC-2017-21845), the Spanish State Research Agency (BCBL “Severo Ochoa” excellence accreditation, SEV-2015-490), the Basque Government (BERC 2018-2021 and PIBA_2019_104), the Spanish Ministry of Science, Innovation and Universities (MICINN; PID2019-105520GB-100 and FJCI-2017-31814), and the Eunice Kennedy Shriver National Institute of Child Health and Human Development of the National Institutes of Health under award number K12HD073945.

## Footnotes

1 https://www.adinstruments.com/support/knowledge-base/it-possible-measure-expired-gasses-partial-pressure-mmhg-rather-percentage. We used the formula *CO*_2_[*mmHg*]=(*P_atm_–P_vap_*)[*mmHg*]·10·*CO*_2_[*V*]/100[*V*], where *CO*_2_[*V*] is the original CO_2_ timeseries, *P_atm_* is the atmospheric pressure in the laboratory at the moment of acquisition, and *P_vap_* is the water vapour pressure associated with expired air. The values of *P_atm_*= 759 and *P_vap_*= 47 were used for all the sessions.

