## Supplementary Material for "ICA-based Denoising Strategies in Breath-Hold Induced Cerebrovascular Reactivity Mapping with Multi Echo BOLD fMRI"

a. Basque Center on Cognition, Brain and Language, Donostia, Spain
b. University of the Basque Country UPV/EHU, Donostia, Spain
c. Scientific and Statistical Computing Core, NIMH/NIH/HHS, Bethesda, MD
d. Physical Therapy and Human Movement Sciences, Feinberg School of Medicine, Northwestern University, Chicago, IL
e. Biomedical Engineering, McCormick School of Engineering, Northwestern University, Evanston, IL

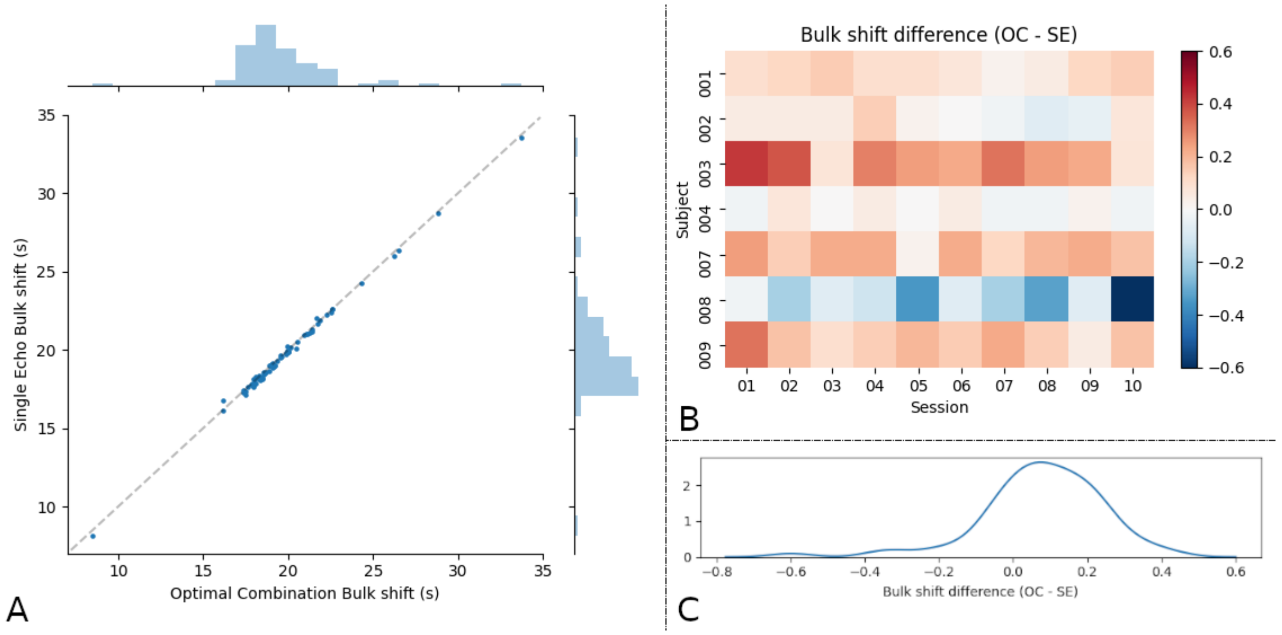


*Supplementary figure 1: (A) Bulk shift of the alignment of the PETCO2 trace with the average GM timeseries of the Optimally Combined (OC) vs Second Echo (SE). The two bulk shifts strongly correlate (r=1, p<0.001). (B) Bulk shift difference of the alignment of the PETCO2 trace with the average GM timeseries of the Optimally Combined (OC) vs Second Echo (SE). The average bulk shift difference across all subjects and sessions is 0.08 ± 0.17, which can be seen as negligible considering the TR of the acquisition (1.5 s) and the shift step used for the lag optimization (0.3 s), since the bulk shift computed from the OC data is not significantly delayed compared to the bulk shift of the SE data (one sample t-test with sample size=7, t = 1.22, p>0.1) (C) Bulk shift difference distribution.*

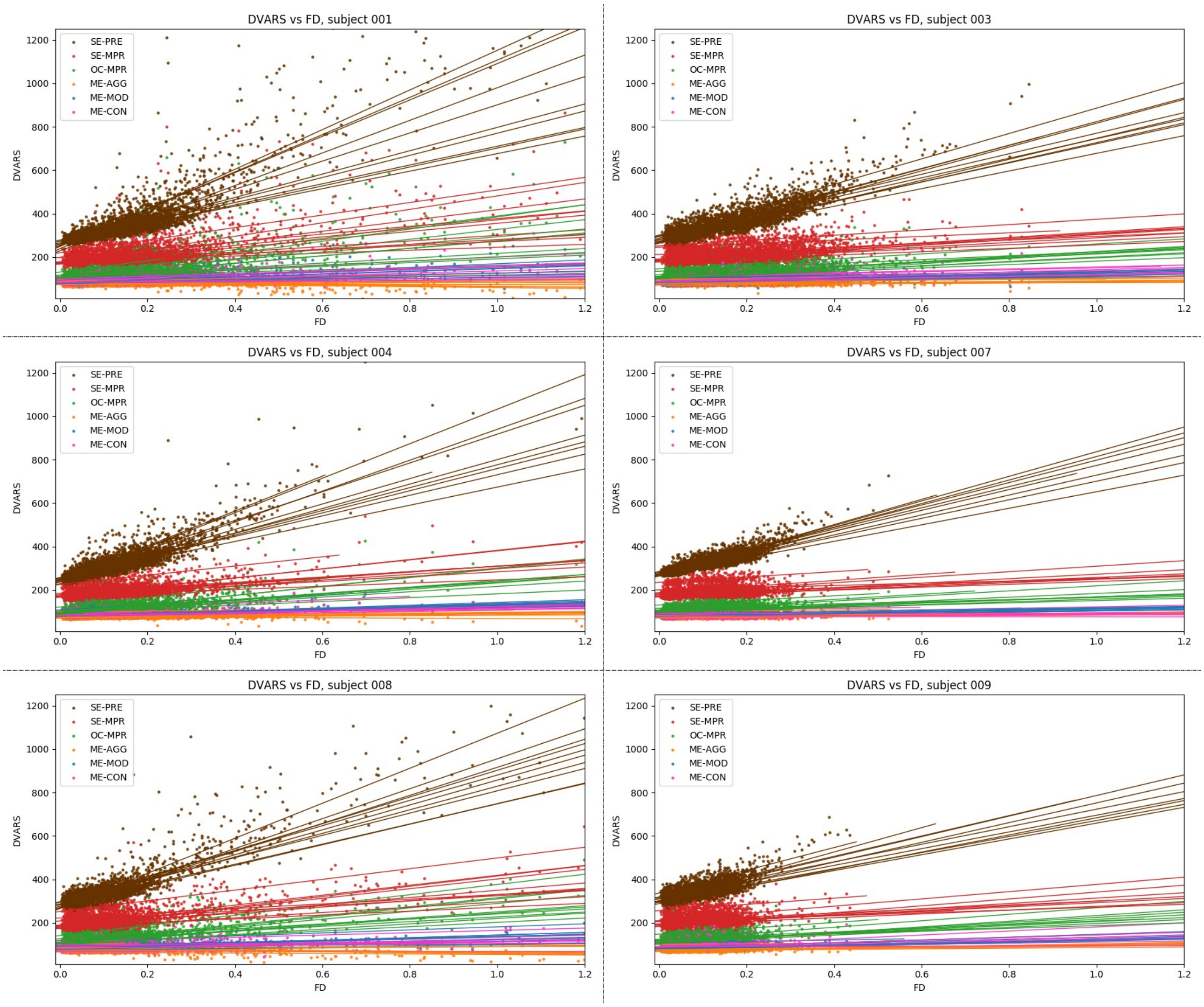


*Supplementary figure 2: Relation between the DVARS of the different analysis pipelines and FD for all the subjects (subject 002 is in the main text). Each point represents a timepoint, each line the linear regression between both timeseries in a session. SE-PRE: raw data; SE-MPR: single-echo; OC-MPR: optimally combined; ME-AGG: aggressive; ME-MOD: moderate; ME-CON: conservative.*

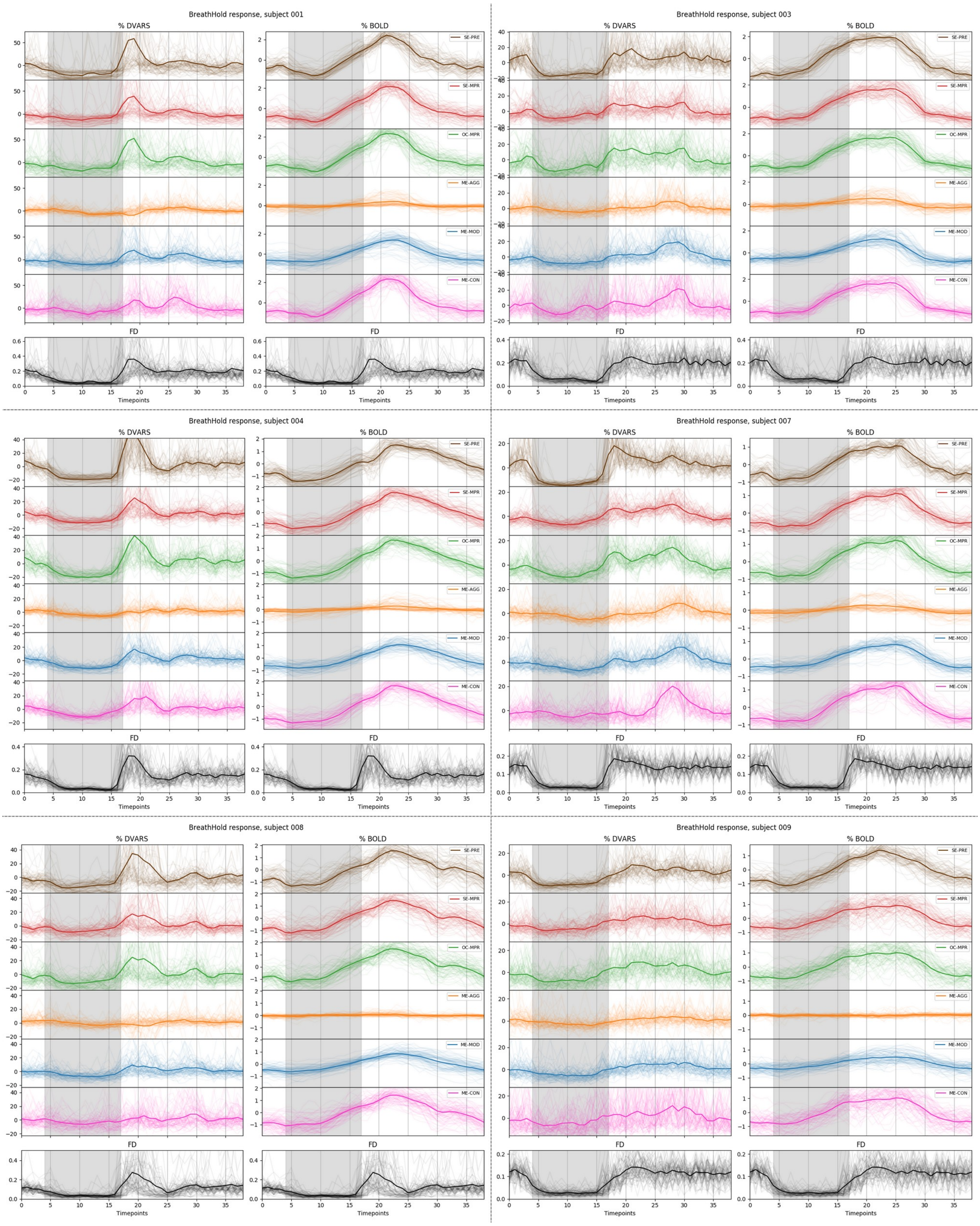


*Supplementary figure 3: Average %DVARS and GM %BOLD response of all BH trials across ten sessions for all subjects (traces for subject 002 can be found in the main text). Each transparent line is a trial, the solid line is the average across all the trials. SE-PRE: raw data; SE-MPR: single-echo; OC-MPR: optimally combined; ME-AGG: aggressive; ME-MOD: moderate; ME-CON: conservative.*

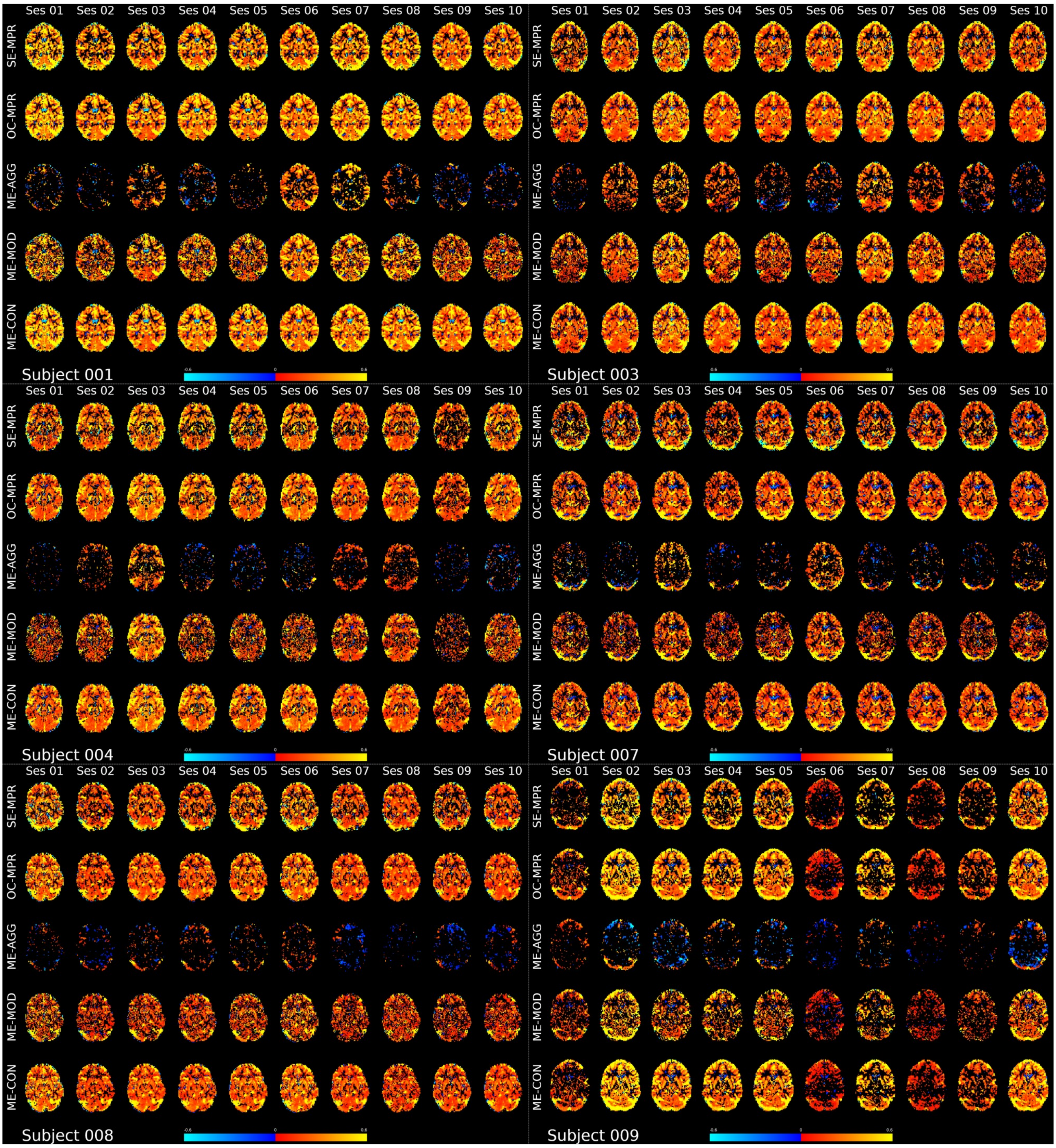


*Supplementary figure 4: Thresholded CVR map obtained with the different lagged-GLM analysis for all sessions of all subjects (maps of subject 002 are available in the main text). SE-PRE: raw data; SE-MPR: single-echo; OC-MPR: optimally combined; ME-AGG: aggressive; ME-MOD: moderate; ME-CON: conservative.*

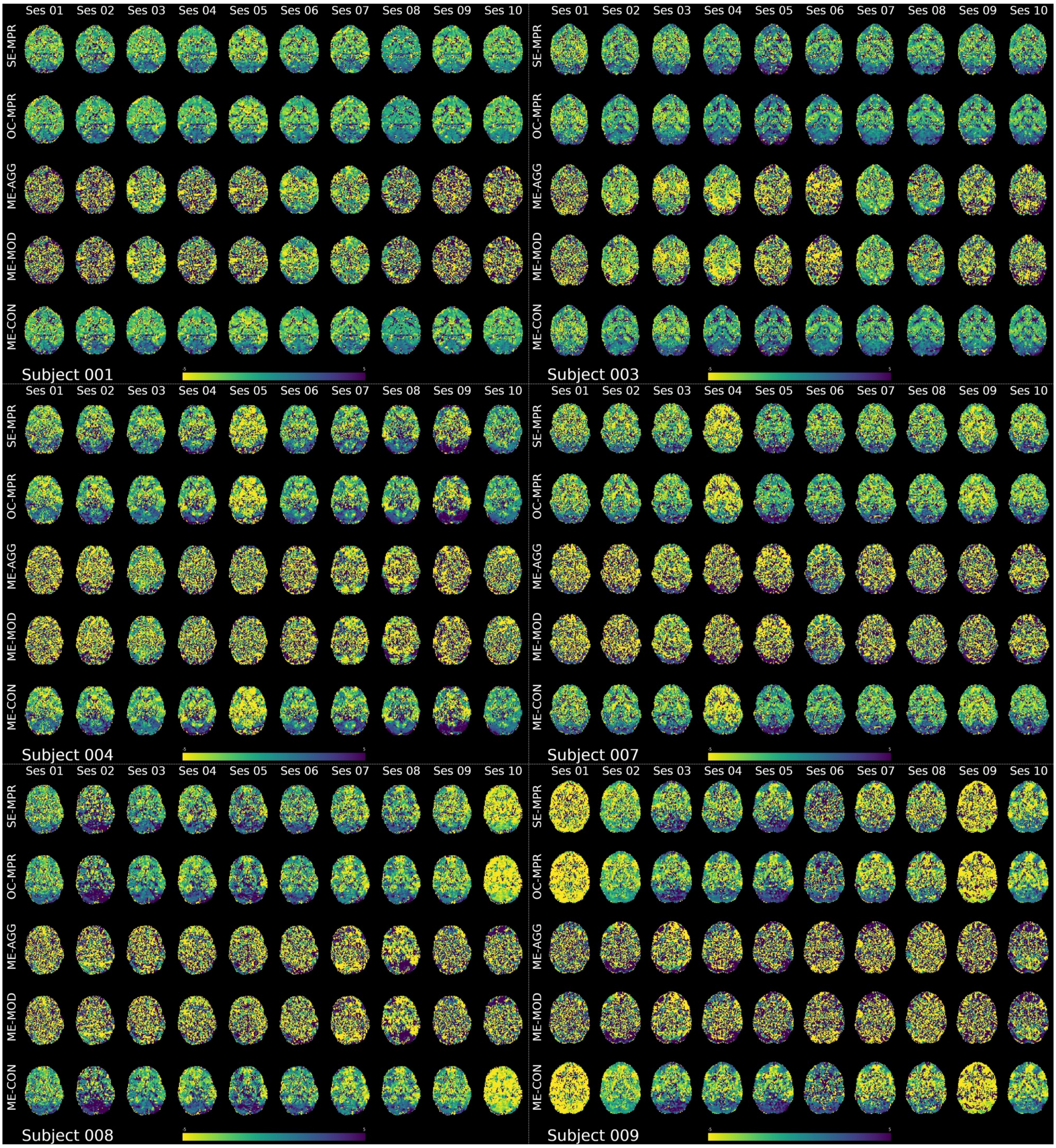


*Supplementary figure 5: Untresholded lag map obtained with the different lagged-GLM analysis for all sessions of all subjects (maps of subject 002 are available in the main text). SE-PRE: raw data; SE-MPR: single-echo; OC-MPR: optimally combined; ME-AGG: aggressive; ME-MOD: moderate; ME-CON: conservative.*

| Subject | Average value | SE-MPR | OC-MPR | ME-AGG | ME-MOD | ME-CON |
|---|---|---|---|---|---|---|
| 001 | CVR [%BOLD/mmHg] | 0.26 | 0.25 | 0.1 | 0.22 | 0.24 |
| | Lag [s] | -0.74 | -0.6 | -0.31 | -0.39 | -0.5 |
| | T value | 7.87 | 9.73 | 1.79 | 12.52 | 10.44 |
| | % significant voxels | 4.41 | 5.26 | 0.95 | 3.83 | 5.39 |
| 002 | CVR [%BOLD/mmHg] | 0.17 | 0.17 | 0.07 | 0.15 | 0.17 |
| | Lag [s] | 0.13 | 0.3 | 0.19 | 0.07 | 0.28 |
| | T value | 7.5 | 9.77 | 1.72 | 11.87 | 10.32 |
| | % significant voxels | 5.76 | 7.08 | 1.04 | 4.79 | 7.18 |
| 003 | CVR [%BOLD/mmHg] | 0.19 | 0.17 | 0.11 | 0.16 | 0.17 |
| | Lag [s] | -0.23 | 0.01 | -0.8 | -0.5 | 0.04 |
| | T value | 6.73 | 8.07 | 3 | 10.8 | 8.44 |
| | % significant voxels | 4.75 | 6.01 | 1.71 | 4.45 | 6.17 |
| 004 | CVR [%BOLD/mmHg] | 0.21 | 0.2 | 0.07 | 0.17 | 0.2 |
| | Lag [s] | -1.31 | -1.3 | -0.98 | -0.98 | -1.25 |
| | T value | 7.09 | 8.75 | 1.94 | 10.6 | 9.42 |
| | % significant voxels | 3.79 | 4.77 | 0.92 | 3.24 | 4.89 |
| 007 | CVR [%BOLD/mmHg] | 0.16 | 0.14 | 0.08 | 0.13 | 0.14 |
| | Lag [s] | -1.03 | -0.83 | 0.28 | -0.5 | -0.75 |
| | T value | 6.29 | 7.56 | 1.54 | 8.46 | 7.94 |
| | % significant voxels | 3.65 | 4.9 | 0.94 | 3.08 | 5 |
| 008 | CVR [%BOLD/mmHg] | 0.18 | 0.17 | 0.03 | 0.14 | 0.16 |
| | Lag [s] | -1.1 | -1.14 | -0.36 | -0.52 | -1.08 |
| | T value | 6.74 | 8.03 | 0.47 | 9.64 | 8.42 |
| | % significant voxels | 4.43 | 5.5 | 0.79 | 3.71 | 5.69 |
| 009 | CVR [%BOLD/mmHg] | 0.2 | 0.19 | -0.02 | 0.16 | 0.18 |
| | Lag [s] | -2.16 | -2.1 | 0.64 | -0.56 | -2.07 |
| | T value | 5.33 | 6.27 | -0.62 | 7.22 | 6.34 |
| | % significant voxels | 2.48 | 3.53 | 0.59 | 2.38 | 3.64 |

*Table 1: Subject average CVR, lag, T value and percentage of statistical voxels in the white matter across strategies (values for grey matter are in the main text). SE-PRE: raw data; SE-MPR: single-echo; OC-MPR: optimally combined; ME-AGG: aggressive; ME-MOD: moderate; ME-CON: conservative.*

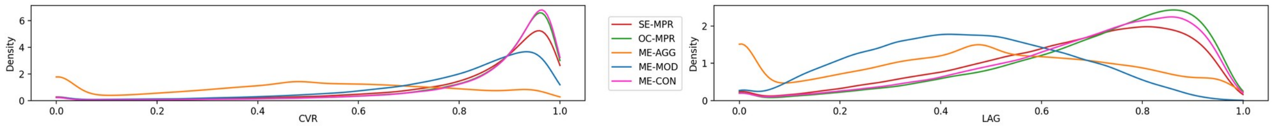


*Supplementary figure 6: Grey matter distribution of ICC scores across voxels for all the pipelines. SE-PRE: raw data; SE-MPR: single-echo; OC-MPR: optimally combined; ME-AGG: aggressive; ME-MOD: moderate; ME-CON: conservative.*
